# Ecological context shapes microbial contributions to nutrition and development in *Drosophila melanogaster*

**DOI:** 10.1101/2025.10.14.682149

**Authors:** Elisabeth K. Riedel, Laura Lender, Vienna Kowallik, Melanie M. Pollierer, Marko Rohlfs

## Abstract

Microbial symbionts play a critical role in shaping insect development by supplying essential nutrients, influencing host phenotype, and mediating context-dependent mutualistic outcomes. In the fruit fly *Drosophila melanogaster*, larvae develop in ephemeral breeding substrates such as decaying fruits, where they acquire maternally transmitted bacterial and fungal symbionts. These substrates vary in their structural and chemical properties, potentially altering the composition and function of associated microbial communities. Here, we combine semi-natural microcosm experiments with compound-specific isotope analysis of essential amino acids to assess how variation in fruit substrate and symbiont origin affect larval nutrition, development, and adult phenotypes. We find that microbial community composition and environmental context jointly shape amino acid flow, with larvae deriving variable proportions of their nutrition from plant, bacterial, and fungal sources. These differences translate into distinct developmental outcomes, including shifts in larval development time – either delayed or accelerated, depending on the symbiont-substrate combination – and significant variation in adult body weight. Our findings introduce symbiont-environment interactions as a framework for understanding how ecological context shapes nutritional mutualisms, showing that environmental heterogeneity can alter microbial composition, nutrient sources, and larval performance. Together, these results reveal how ecological context modulates mutualistic dependence and underscores the context sensitivity of host-microbe associations.

## Introduction

The evolutionary success of insects in occupying a wide variety of ecological niches is outstanding (1). Interactions with microorganisms, such as bacteria and fungi, which lead to the formation of temporarily stable or transient host-microbe symbioses contribute to this success (2,3). Symbiotic microbiota facilitate the realization of the host’s ecological niche through nutritional supplementation, protection from pathogens, and detoxification or other environmental modifications (4–8). The composition of symbiotic microbiota is shaped by a diversity of both microbe-microbe interactions and host control mechanisms, which operate across open ecosystems to precise genetic shaping and vertical transmission (9).

In systems where both hosts and microbial symbionts can associate flexibly with multiple partners (10), ecological context becomes a key driver of symbiont community composition, with potentially significant consequences for host phenotypes and the stability of the mutualism (11). This is especially true for the vast diversity of insects that develop in ephemeral resource patches, such as dung, carrion, mushroom, or rotting plant tissue (12). The benefits of microbiota that vary across time and space are less evident than those of mutualistic partners with well-defined functions or identities (10,13–15). Rather than symbiont composition or environmental factors alone, it is their dynamic interplay that ultimately determines the expression of insect life-history traits and, by extension, the breadth of the host’s ecological niche (10).

Cases where the impact of microbial symbionts on host development depends on the breeding environment, defined as symbiont-environment interactions, remain largely unexplored regarding the arising benefits and costs for the host (15). In this study, we address this gap by investigating how variation in the breeding environment influences symbiont community composition, nutrient acquisition during larval development, and the resulting host phenotype in the fruit fly *Drosophila melanogaster*.

*Drosophila melanogaster*, a cosmopolitan decomposer insect, is a representative of the vast diversity of insect species that are generalists for transient breeding sites, including dung, carrion, fruits, and plant litter. For example, *D. melanogaster* was observed to breed in 15 of 24 plant substrates, primarily fruits, examined by Atkinson and Shorrocks (16). During juvenile development the larval stages of *Drosophila* spp. engage in prolonged and intimate interactions with microbiota residing both inside their gut and in the decaying plant substrate (17,18). Symbionts serve as a vital source of amino acids within predominantly low-protein plant substrates (19,20) thus, playing a pivotal role in shaping the expression of adult life-history traits that are largely contingent upon preadult growth and development (21). Furthermore, dietary symbionts can also exhibit a protective role against pathogens like toxin-producing filamentous mold fungi in *Drosophila* larvae (22). The deposition of fecal maternal microbiota during egg-laying significantly influences the symbiont community composition in localized breeding patches (23–27). Additionally, the inherent characteristics of the breeding substrate and seasonal variation seem to have a profound influence on the symbiont composition of *Drosophila* (14,28–30). Beyond seasonal weather variation, a complex interplay of factors, including plant substrate properties such as pH, water availability, and nutrient content, along with microbial interactions, shapes these dynamics (31). To date, how variation in symbiotic microbes influence *D. melanogaster* development across different environments and thus contribute to generalization in mutualism (10) remains an open question. In particular, the consequences of fruit substrate variation, and its impact on the bacterial and fungal symbionts that provide essential nutrition during larval development, is unknown.

Carbon (^13^C) and nitrogen (^15^N) stable isotope analysis offers an effective method for uncovering the specific plant substrates that serve as nutritional sources for *Drosophila* larvae (32). However, bulk stable isotope analyses fall short in distinguishing between plants and microorganisms as nutritional sources for fruit fly larvae. Compound specific stable isotope analysis (CSIA) of essential amino acids (EAA) is an emerging tool to investigate both basal resources and trophic positions of consumers (33–36). Unique patterns, or ‘fingerprints’, of δ^13^C in EAA produced by bacteria, fungi or plants allow to quantitatively trace the biosynthetic origin of these EAA in consumers (36–38). We used this approach to quantify, for the first time, how plant substrates, bacteria, and fungi differentially contribute to larval nutrition in *Drosophila*, depending on breeding environment and symbiont composition.

We analyzed EAA profiles of adult *Drosophila* and the respective ontogenetic environment. These adult flies developed from larvae reared under distinct symbiont–environment combinations, established by inoculating different plant substrates with female-derived fecal microbiota from flies maintained in source microcosms. The microcosms, based on field-exposed substrates, sustained fly-symbiont associations and enabled successive microbial transmission across generations. Generally, we used two types of microbiota-inoculated plant substrates: substrate with symbionts from females that lived in the same substrate environment (autochthonous symbiont inoculum, *autoch*SI) and substrate with symbionts from females that came from a different substrate environment (allochthonous symbiont inoculum, *alloch*SI; Fig 1). We employed a metabarcoding approach to, first, characterize the bacterial and fungal symbionts transmitted through fecal matter deposition by females from different substrate environments (39). Microdiversity of strain diverse bacterial phylotypes was assessed to reveal the potential of environment-specific symbiont strain selection (40). Second, we analyzed microbiota changes during *Drosophila* development in substrate with *autoch*SI and *alloch*SI. Lastly, we recorded larval development time and biomass gain through quantification of adult body weight to investigate how the consequences of *autoch*SI versus *alloch*SI are connected to larval nutrient intake and variation in the expression of insect life-history traits. In this context, it is assumed that ephemeral resources, which can be subject to unpredictable catastrophic events (12) exert selective pressure for rapid development while simultaneously selecting for maximal biomass gain as key fitness components in holometabolous insects (21,41).

**Fig 1.**
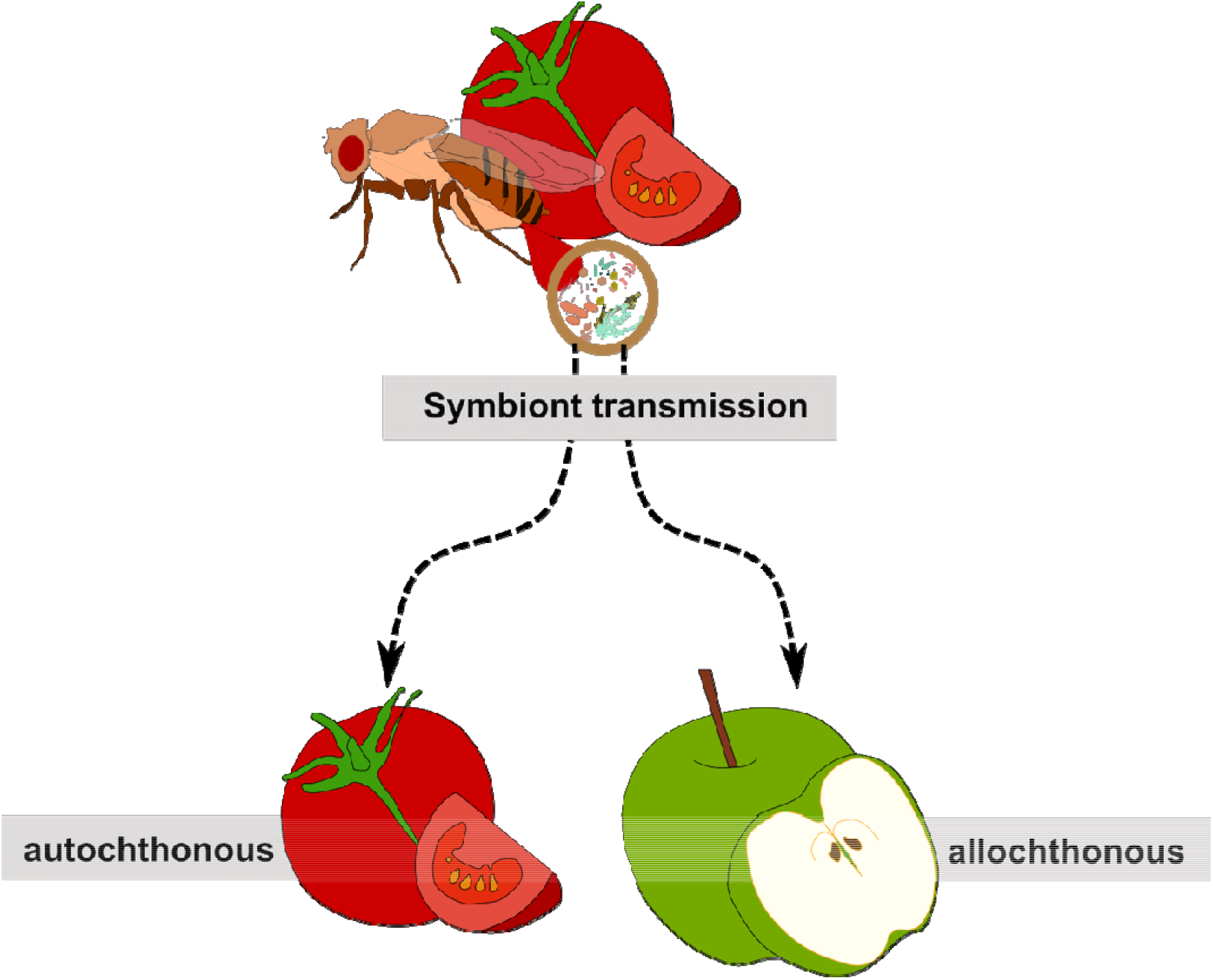
Ecological context as determined by autochthonous versus allochthonous symbiont transmission may cause variation in the expression of insect life-history traits. Many insects develop in short-lived, ephemeral resource patches. The subsequent generation of adult insects may encounter similar resource patches and reproduce there. However, due to the temporal and spatial variability in the occurrence of these habitats, dispersing individuals may instead encounter novel and dissimilar breeding sites that differ substantially in both structure and chemical composition. Microbial symbionts, which are essential for larval development and transmitted maternally, are thereby exposed to new environmental conditions. These shifts may alter symbiont growth dynamics and community composition, potentially affecting host developmental outcomes. Thus, the breeding substrate type determines whether symbionts supporting juvenile development encounter a familiar (autochthonous) or a novel (allochthonous) ecological context.

By emphasizing the often overlooked yet pervasive influence of environmental heterogeneity on symbiont composition in a generalist host-microbe system, our study advances the understanding of nutritional symbioses shaping host development and mutualistic dependence.

Thus, it provides new insights into the limitations that environmental heterogeneity imposes on niche width (42,43) and the evolution of mutualistic symbioses in animal hosts that have been selected to reproduce in ephemeral resource patches (12,44,45). More broadly, we propose symbiont-environment interaction as a conceptual framework for uncovering the ecological processes that generate variation in partner breadth and mutualistic dependence (10,11).

## Material and methods

### *Drosophila* stock and rearing

The *Drosophila melanogaster* laboratory stock originated from wild-caught individuals collected in Kiel, Germany in 2006 (46). Since then, the population has been maintained as an outbred stock of several hundred individuals per generation. Flies are reared on a standard laboratory diet (S1 Text), under controlled conditions of 20 °C and a 16:8 h light-dark cycle.

### Field-derived microbes and substrate-specific *Drosophila* microcosms

*Drosophila melanogaster* reared in laboratory environments typically harbor a reduced and less diverse microbiota compared to their wild counterparts (18,28,30). To generate microbiota inocula more representative of natural conditions, we aimed to recolonize our laboratory *D. melanogaster* population with field-acquired microbial symbionts. In September 2019 we exposed cut apple and tomato substrates on enclosed pedestals coated with tangle-trap paste (TEMMEN Insektenleim, Temmen GmbH) allowing access to flying insects only. This exposure allowed the substrates to be colonized by environmental microbes and insects visiting the patches. After one week, we retrieved the baits and removed any remaining arthropods. The microbially enriched substrates were then provided as the sole food source to our laboratory stock of *D. melanogaster*, thereby introducing field-derived microbiota to the lab-reared flies. After 24 hours, these flies were transferred to fresh apple or tomato pieces to facilitate egg laying and the transmission of associated microbes. New fly populations were subsequently reared from these substrates, with the respective fruit type regularly replenished to maintain continuous development and microbial transfer across generations. *Drosophila*-microbe microcosms were maintained in 22 L (39 × 28 × 28 cm) plastic containers, where adult flies had continuous access to water and a mixture of sucrose and hydrolyzed brewer’s yeast. Approximately once per month, microcosms were renewed by allowing flies to oviposit into fresh pieces of the respective fruit substrate, which were then transferred to a new container. The number of these transfer events was recorded for each microcosm. Female flies from the populations were used as symbiont donors and are referred to as “apple” and “tomato” flies. A second set of “apple” and “tomato” populations was created in a similar manner in 2020.

### Collection of female fecal symbionts

A fundamental assumption of this study is that bacterial and fungal microbiota transmitted via female fecal deposition play a major role in establishing the larval microbiota and shaping offspring-microbe symbioses during development (23–27). To minimize the effects of strong inter-individual variation in microbial transmission (25), we standardized the deposition of female-derived symbionts across all microbiota community analyses and experiments described herein. To standardize symbiont inoculation, 30 female flies from each tomato- or apple-associated population were placed in sterile 15 mL screw-cap tubes and allowed to defecate overnight, depositing their microbiota as fecal droplets on the tube walls. For metabarcoding, flies were removed, and the tubes were immediately frozen at −80 °C until DNA extraction (see below). For experiments assessing essential amino acid flow and larval development, freshly collected fecal material served as the inoculum. In these cases, after removing the flies, the fecal deposits were suspended in 5 mL phosphate-buffered saline and dislodged by shaking (180 rpm) for 60 minutes. Aliquots of 50 µL were pipetted into each experimental unit (see below).

Microbiota were collected from the 2019 and 2020 microcosms after varying numbers of transfer events. Details of the microcosms used in the larval development experiments are provided in S2 Table. For metabarcoding-based characterization of symbiont communities from “apple” and “tomato” flies, we sampled fecal material from both apple and tomato microcosms established in 2019 after twelve and seventeen transfer events, and from the 2020 microcosms after one, two, and five transfer events. Fecal material from apple-associated flies experimentally applied to apple substrate is termed an autochthonous symbiont inoculum (*autoch*SI), whereas the same material on tomato substrate is termed allochthonous (*alloch*SI). Conversely, fecal material from tomato-associated flies is *autoch*SI on tomato and *alloch*SI on apple (Fig 1).

### Metabarcoding of *Drosophila melanogaster* bacterial and fungal microbiota

#### DNA-Extraction and amplicon sequencing

For the extraction of microbial DNA from fly fecal material, we adapted the protocol described by Fink et al. (47). Briefly, screw-cap tubes containing frozen samples were thawed, and fecal deposits adhering to the tube walls were collected using sterile swabs pre-moistened with sterile phosphate-buffered saline (PBS). The cotton tip of each swab was aseptically removed with sterilized scissors and placed directly into the first reaction tube of the Qiagen DNeasy PowerSoil Kit. To each sample, we added 60 μL of the kit’s buffer C1 along with 20 μL of proteinase K, followed by incubation for 2 h at 50 °C in a thermomixer to facilitate cell lysis. Samples were then vortexed for 10 min using a vortex adapter before proceeding with the manufacturer’s standard protocol. Two empty screw-cap tubes, processed alongside the fecal samples, served as negative controls for the extraction workflow. For the analysis of symbiont communities in the developmental substrate (see substrate preparation below in *Larval development experiment*), we used material collected after the development of single larvae in the same experimental setup described for the larval development assays (see below). Surface scrapings from substrates were randomly pooled in groups of 10 experimental units, and DNA was extracted using the Qiagen DNeasy PowerSoil Kit according to the manufacturer’s protocol.

The DNA samples were sent to Advanced Identification Methods (AIM) GmbH, Leipzig, Germany, for dual-tag library preparation, amplicon sequencing, and demultiplexing. For bacterial community profiling, the V3–V4 hypervariable region of the 16S ribosomal RNA gene using primers 341F and 785R (48) was amplified. Fungal communities were profiled by amplifying the internal transcribed spacer 2 (ITS2) region with primers ITS3_KYO2 (forward) and ITS4_KY03 (reverse) (49), which are optimized for broad taxonomic coverage of fungi. In total, eleven fecal samples, nine substrate-scrape samples, and two negative extraction controls were processed. Libraries were sequenced on an Illumina MiSeq platform (v3 chemistry) using a paired-end strategy (2 × 300 bp), generating high-resolution reads suitable for downstream community composition analyses.

#### Taxonomic identification

The demultiplexed FASTQ files were processed in QIIME 2 version 2022.2 (50) on a Linux-based virtual machine (kernel 5.4.0-148-generic). Sequence quality control and denoising were performed using the *denoise-paired* command, which implements the DADA2 algorithm (51) as integrated within the QIIME 2 framework, using default settings unless stated otherwise below.

For the 16S sequences, forward reads were truncated at 299 bp and reverse reads at 255 bp after visual inspection. Primer sequences were trimmed from the start of each read (17 bp from forward and 21 bp from reverse), leaving an overlap of ∼110 bp for merging. Taxonomic assignment was performed in QIIME 2 using the q2-feature-classifier plugin (52) with a naïve Bayes classifier. The classifier was trained against the SILVA database release 132 April 2018 (53), reduced to 99% identity sequences with taxonomy resolved to seven levels limited to sequences containing our specific primer set.

Due to length variability in the ITS2 region, which increases the likelihood of primer read-through, primer sequences and their reverse complements were removed using *Cutadapt* in QIIME 2 (54). Quality trimming involved removing the first 30 low-quality bases from both forward and reverse reads, and truncating reverse reads at 215 bp after visual quality inspection. Forward reads were processed with a quality threshold of 2.0, while reverse reads were allowed an error rate of up to 5.0 due to lower overall quality. We did not filter for chimeras since they are much less likely to occur when using ITS as a marker gene (55). Taxonomic assignment was based on the UNITE database for fungal sequences, adapted for the QIIME 2 environment. We used the “dynamic” taxonomy, in which clustering thresholds vary between 1–3% depending on fungal lineage characteristics (56). A naïve Bayes classifier was trained on the full taxonomy, following QIIME 2 development team recommendations (57), and applied to our ITS reads for taxonomic identification (52). A total of 136 unassigned amplicon sequence variants (ASV) were manually blasted (BLAST+ 2.13.0, December 2022) against the NCBI “Nucleotide collection (nr/nt),” including GenBank, EMBL, DDBJ, PDB, and RefSeq. Most unassigned features were of non-fungal origin (plants, animals, etc.) and were excluded from further analysis. Three additional fungal ASVs were identified and manually added to the taxonomy table: one Saccharomycetaceae species (one read, NCBI accession: 7ZW0_LD, 100% coverage) and two features belonging to the *Penicillium chrysogenum* complex (two reads each, NCBI accession: MW774585.1, 98.39% coverage).

#### Microbial community analysis

All analyses of microbial alpha and beta diversity, as well as taxonomic composition at the genus, species, and ASV levels, were performed in R (R Core Team, 2022; RStudio Team, 2022) using the *phyloseq* package (58). For fungal community analysis, ASVs represented by a single read were removed as likely artifacts, and samples with fewer than 1,000 reads were excluded. This resulted in the removal of one pooled tomato substrate sample. Potential contaminants were identified using the *decontam* package (59) with a threshold-based selection strategy. After filtering, the dataset contained 2,051 taxa represented by more than 1.1 million reads across 16 samples. Similarly, for bacterial communities we used the *decontam* package to identify and remove 27 putative contaminants based on prevalence. Blank samples, empty taxa, and sequences assigned to chloroplasts or mitochondria were excluded. ASVs represented by a single read were removed, and one apple substrate sample was excluded due to low read counts (<1,000). The final dataset comprised 614 taxa represented by ∼300,000 reads across 16 samples. Rarefaction curves indicated sufficient sequencing depth for all samples, with asymptotic behavior reached in each case (Fig S1). Therefore, no rarefaction was applied, and analyses proceeded with the full dataset.

*Alpha* and *beta* diversity of fungal and bacterial communities were analyzed separately for substrate and fecal microbiota. Alpha diversity was quantified as Shannon diversity and observed richness (taxon count) using the *estimate_richness* function in the *phyloseq* package. Richness estimates were compared using Wilcoxon rank-sum tests (*stats* package). *Beta* diversity was calculated as Bray-Curtis dissimilarity (60) and visualized by principal coordinates analysis (PCoA) using multidimensional scaling (MDS). Differences in community composition were tested by permutational multivariate analysis of variance (PERMANOVA) with the *adonis* function in the *vegan* package (61).

To assess strain-level variation in the fecal microbiota and its potential ecological relevance, we rarefied samples to an even sequencing depth of 9,400 reads, matching the smallest sample size. We selected three phylotypes identified to species level that contained at least five unique ASVs within each sample type, *Klebsiella pneumoniae, Bacillus subtilis*, and *Lactobacillus buchneri*, as well as *Caulobacteraceae* sp., which reached high relative abundance in fecal microbiota from apple environments (up to 80% of bacterial reads; Fig 2). These taxa were analyzed for *alpha* and *beta* diversity at the strain (ASV) level, following the procedures described above, to explore whether substrate-dependent environments are associated with fine-scale genetic variation in dominant symbionts.

**Fig 2.**
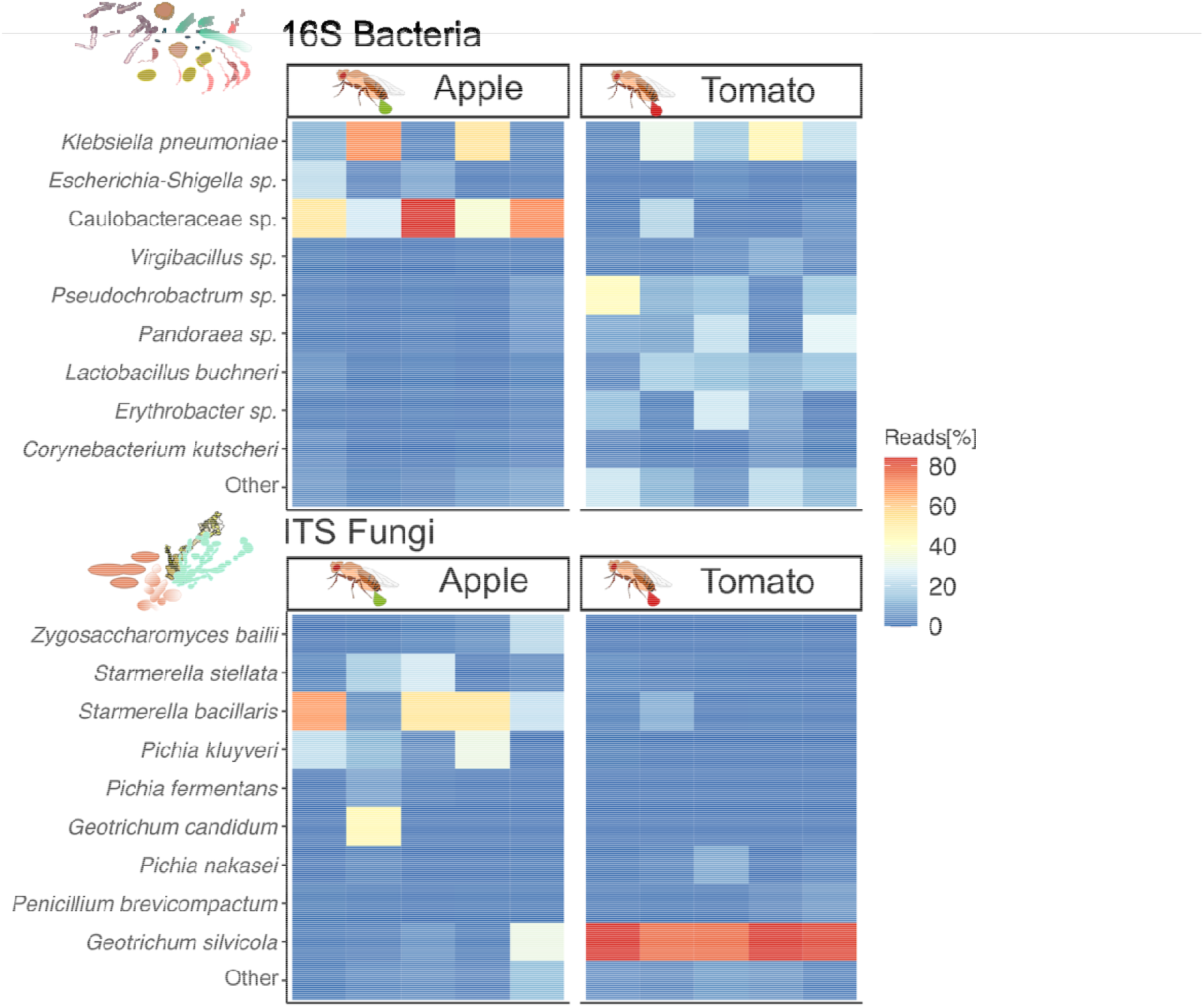
Breeding environment determines *Drosophila melanogaster* symbiont community composition. Heatmaps showing the nine most abundant bacterial and fungal taxa found in the fecal material of *Drosophila melanogaster* females from apple and tomato environments (n = 5). The ITS2 region and the 16S ribosomal subunit were used as marker genes for identification of fungal and bacterial communities. Amplicon sequence variants were agglomerated at species level and the nine most abundant taxa are displayed. All other taxa are grouped under “Other”. The relative abundance of taxa in each sample is depicted as color intensity from blue to red as percentage of total reads.

### Compound specific isotope analysis of amino acids

Compound-specific isotope analysis (CSIA) of essential amino acids (EAAs) was performed using fly and substrate material from the October 2021 run of the larval development assay (see below). Within each treatment, fly samples were pooled by stratified random sampling (n = 8 per pool) to obtain at least 2 mg dry weight (following Larsen et al. 2013), resulting in five replicate pools per treatment. Developmental substrates from these assays were frozen, pre-dried for 24 h at 65 °C, lyophilized, and pooled accordingly. Fresh substrate samples were also dried (48 h at 65 °C) and lyophilized. For each substrate type (apple, tomato, and additionally, standard laboratory diet), five replicates (10-15 mg dry weight) were prepared for amino acid extraction. Flies and substrates were processed in separate extraction batches for synchronized handling, with samples randomized across treatments.

Amino acids were extracted and derivarized as described in Larsen et al. (62). Derivatized amino acids were analyzed in triplicate by gas chromatography–combustion isotope ratio mass spectrometry (GC-C-IRMS). They were injected into a Thermo Finnigan Trace 1310 gas chromatograph coupled to a Delta Plus IRMS (Finnigan, Bremen, Germany), equipped with an Agilent J&W VF-35ms GC column (30 m × 0.32 mm × 1.00 µm). Carbon isotope values were corrected for derivatization (63) and expressed relative to Vienna Pee Dee Belemnite in δ-notation (‰ ). We obtained isotope values of 12 amino acids including alanine (*Ala)*, asparagine/aspartic acid (*Asx)*, glutamine/glutamic acid (*Glx)*, isoleucine (*Ile*\*), leucine (*Leu*\*), methionine (*Met*\*), phenylalanine (*Phe*\*), proline (Pro), serine (Ser), threonine (*Thr*\*), tyrosine (Tyr) and valine (*Val*\*); asterisks denote essential amino acids.

A linear discriminant analysis (LDA) was trained with δ^13^C values of essential amino acids from bacteria, fungi, and plants, published by Larsen et al. (38,62) and Pollierer et al. (64) to represent potential basal resources. Decision boundaries were constructed to project substrate, substrate post-eclosion, and fly samples into the LDA space, based on their essential amino acid profiles. To estimate the proportional contribution of basal resources used by flies we ran Bayesian mixing models implemented in R package MixSIAR (65) based on mean-centered δ^13^C values of essential amino acids in the potential basal resources as used in LDA and in the flies.

### Larval development experiments

To ensure that variation in larval developmental success could be attributed solely to differences in substrate and symbiont inoculum conditions, all experiments were conducted with axenic larvae. *D. melanogaster* eggs were obtained overnight from the laboratory stock maintained on standard laboratory diet. The following morning, eggs were surface sterilized by dechorionation in 6% sodium hypochlorite for 10 min, followed by repeated rinses, first with ethanol and then with autoclaved tap water. Embryos were transferred to sterile glucose agar plates and incubated at 25 °C for 24 h to allow hatching. Newly emerged first-instar larvae were then used in the development assays. Smear-plating of freshly hatched larvae showed no visible microbial growth on either bacterial- or fungal-selective agar (S1 Text for media composition). Because all larvae originated from the same large, outbred laboratory population, there was no prior adaptation to either substrate or inoculum. Any observed interaction effects would therefore reflect differences in how microbial communities from different origins function within distinct substrate environments, rather than genetic or epigenetic differences among flies.

Across five independent experimental runs (March 2020, September 2020, March 2021, June 2021, and October 2021), we tested the effects of fecal microbial inocula from “apple” and “tomato” flies, transferred to either autochthonous or allochthonous fruit substrates (see above). Larval development was quantified in four of these runs, while the June 2021 run was dedicated exclusively to substrate microbiota analysis. The October 2021 run additionally included amino acid analysis (S1 Table for sample details).

Organic apples (*Holsteiner Cox*) and vine tomatoes were purchased from a local store (ALECO, Bremen). Apples were peeled and cored, while seeds were removed from tomatoes. Fruits were homogenized to produce a semi-liquid substrate, which was microwaved (800 W) three times until boiling commenced. This treatment standardized substrate texture and moisture content while substantially reducing the initial microbial load prior to inoculation. Plating of the prepared substrates showed no visible microbial growth on either bacterial- or fungal-selective agar.

For each experimental unit, 1 mL of prepared fruit substrate was pipetted into a sterile 2 mL collection tube, followed by 50 μL of fecal suspension originating from “apple” or “tomato” flies. A single axenic larva was then transferred into each tube, and the tubes were sealed with autoclaved dental cotton rolls. Experimental units were placed in racks and incubated under the same environmental conditions as the laboratory stock and microcosms (see above). The following day, all units were checked for the presence of alive larva to ensure that only replicates unaffected by accidental mortality during transfer were included in subsequent analyses. The total number of prepared replicates for each run, the origin of the fecal microbiota, and the final number used in statistical analyses are provided in S1 Table. Replicates that passed this quality check were monitored daily to record the number of days from larval transfer to adult emergence. Newly emerged flies were sexed, immediately frozen, and dried over silica gel before weighing on a precision scale. The June 2021 run, designed to generate samples for metabarcoding of the developmental substrate, used only female fecal microbiota from “tomato” flies to establish *autoch*SI or *alloch*SI conditions; developmental parameters from this assay were therefore excluded from analysis. In the October 2021 run, we included an additional treatment with standard laboratory diet without microbial inoculation, enabling comparison of essential amino acid profiles between flies reared on fruit substrates and those reared on the laboratory diet. Larval development on the standard laboratory diet was not further analyzed.

#### Statistical analysis

Our statistical approach was designed to test the hypothesis that host developmental traits are shaped by symbiont–environment interactions, i.e., that the effect of microbial inoculum origin (*autoch*SI vs. *alloch*SI) on *Drosophila* development depends on the substrate in which larvae develop. Interaction terms between substrate type, inoculum origin, and sex were therefore explicitly included in all models, as significant interaction effects would indicate the presence of such context-dependent symbiont effects.

Model fit was first assessed using data from the initial experimental run for both adult weight and development time. For both traits, a full model including all interaction terms provided an adequate description of the data. We then analyzed data from all four experimental runs, with “run” initially included as a random effect to account for potential variation among runs. For adult weight, a linear mixed-effects model was fitted first. The intraclass correlation coefficient for “run” was low (ICC = 0.03), indicating that between-run variance was negligible. We therefore proceeded with a standard linear model for the final analysis, using z-transformed weight data. The final model structure was:

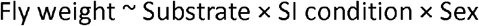

For development time, data were Box–Cox transformed to meet model assumptions and analyzed using a linear mixed-effects model using *lmerTest* (66) with the same fixed-effects structure, yet experimental run was included as a random effect (ICC = 0.45):

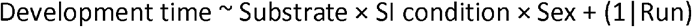

Across all treatments, sample sizes were *n* = 817 for development time and *n* = 796 for weight (exact numbers by treatment are given in S1 Table). Variation in replicate numbers arose from larval mortality and occasional measurement failures. Ninety-five percent confidence intervals for predicted means were calculated from fitted model values using the *predictInterval* function in the *merTools* package (67), based on 1,000 simulated datasets. All statistical analyses of the development assays were conducted in R version 4.2.2 (R Core Team, 2022) using the RStudio interface (version 2022.07.02+576; RStudio Team, 2022).

### Figure design

Graphical outputs involving data from either metabarcoding, CSIA or larval development experiments were generated with the ggplot2 package (68) and adapted in Inkscape v1.1.0 using original designs. Fig 1 was designed in Inkscape. Accessibility of graphics was assessed using Color Oracle 1.3.0.

## Results

### Symbiotic microbiota in the feces of fly populations differ between breeding environments

To assess whether the symbiotic microbiota transmitted by female *D. melanogaster* to egg-laying sites differ by breeding environment, we analyzed fecal material from flies reared in separate populations maintained on distinct substrates (apple or tomato). Across both substrate environments, the fecal material from “apple” and “tomato” flies contained seven bacterial families that comprised at least 80% of the total bacterial microbiota. These families were Bacillaceae, Burkholderiaceae, Caulobacteraceae, Enterobacteriaceae, Lactobacillaceae, Rhizobiaceae, and Sphingomonadaceae. The predominant (at least 70% of all samples) fungal members of the microbiota were yeasts from the order Saccharomycetales (specifically *Pichia* and *Starmerella* species), as well as some *Geotrichum* and filamentous *Penicillium* species (Fig 2, Fig S2). The *alpha* diversity of bacteria showed only marginal difference between symbionts derived from apple and tomato flies (Fig 3A); however, this could be due to between-replicate variability in phylotype numbers (Fig S3). Notably higher diversity among fungal symbionts was observed in the apple flies compared to the tomato flies (Fig 3B, Fig S3).

**Fig 3.**
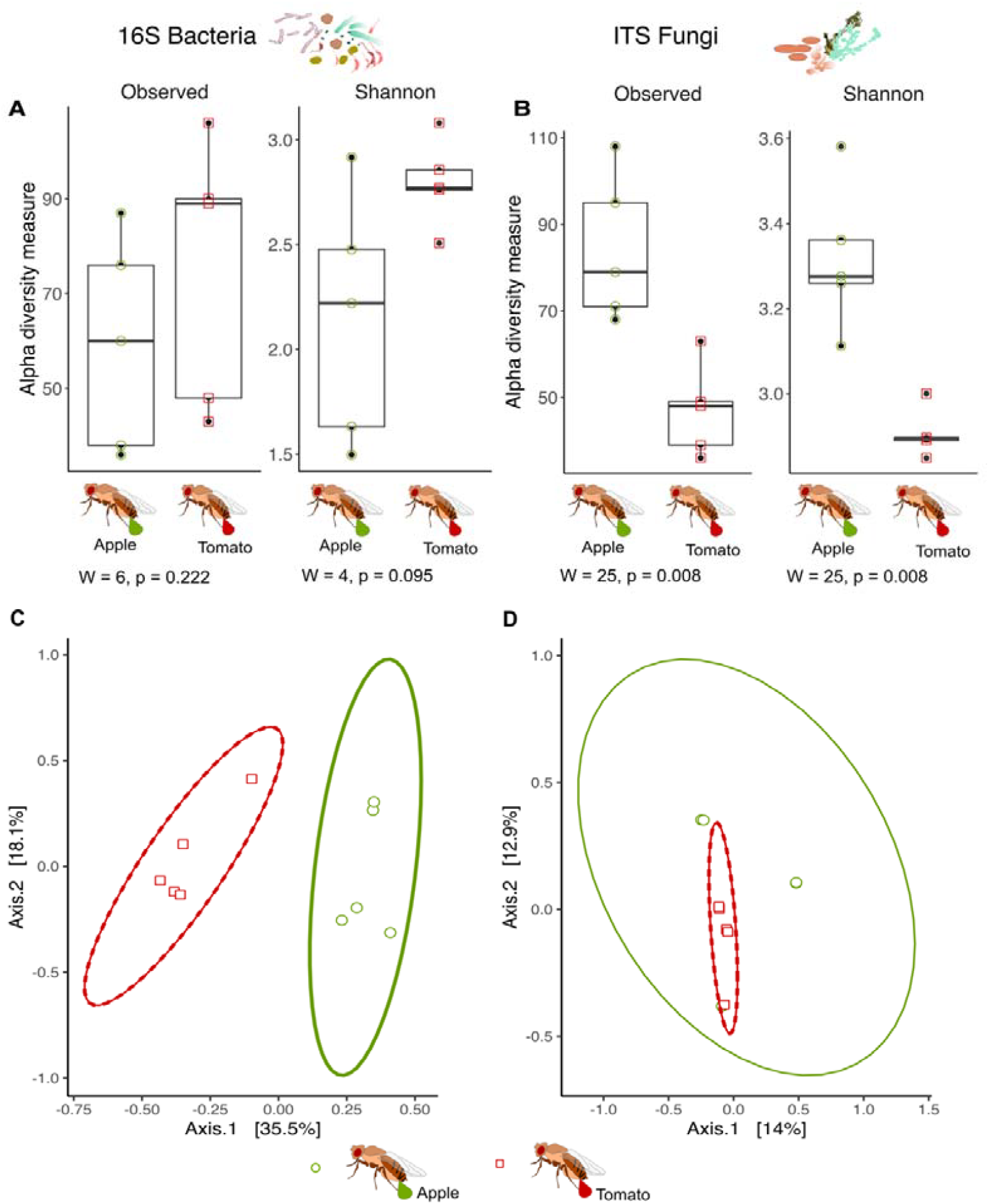
Community composition of *Drosophila melanogaster* symbionts differs between breeding environments. Boxplots depicting observed and Shannon diversity of symbiont communities in the fecal material of *Drosophila melanogaster* females from apple or tomato environments (n = 5). The ITS2 region and the 16S ribosomal subunit were used as marker genes to characterize fungal (A) and bacterial (B) communities, respectively. Symbiont community differences between apple and tomato were assessed using the Wilcoxon Rank-Sum Test. Principal Coordinate Analysis (PCoA) based on Bray-Curtis dissimilarity showing the bacterial (C) and fungal (D) symbiont community composition. Ellipses represent the 95% confidence interval.

The composition of bacterial symbionts differed significantly between the substrate environments (PERMANOVA; F_1,8_ = 4.03, p = 0.012, Fig 3C). Bacterial symbionts from “apple” flies were dominated by a species from the gram-negative Caulobacteraceae family (Fig 2). Also, the fungal symbiont composition exhibited distinctions between “apple” and “tomato” flies, although the differentiation was subtle (PERMANOVA; F_1,8_ = 1.09, p = 0.156, Fig 3D). Notably, symbionts in the tomato environment were predominantly (70-80% of reads) composed of *Geotrichum silvicola* (Fig 2, Fig S2). Additionally, many phylotypes were found to consist of more than five strain variants (S2 Table). A deeper analysis of ASV microdiversity within the most divergent bacterial phylotypes revealed substrate-dependent differentiation in a Caulobacteraceae phylotype, *Klebsiella pneumoniae, Bacillus subtilis*, and the *Lactobacillus buchneri* phylotype (Fig 4). This indicates that environment-driven differentiation in *Drosophila* symbionts is evident not only at the phylotype level but also at the level of intraspecific variation.

**Fig 4.**
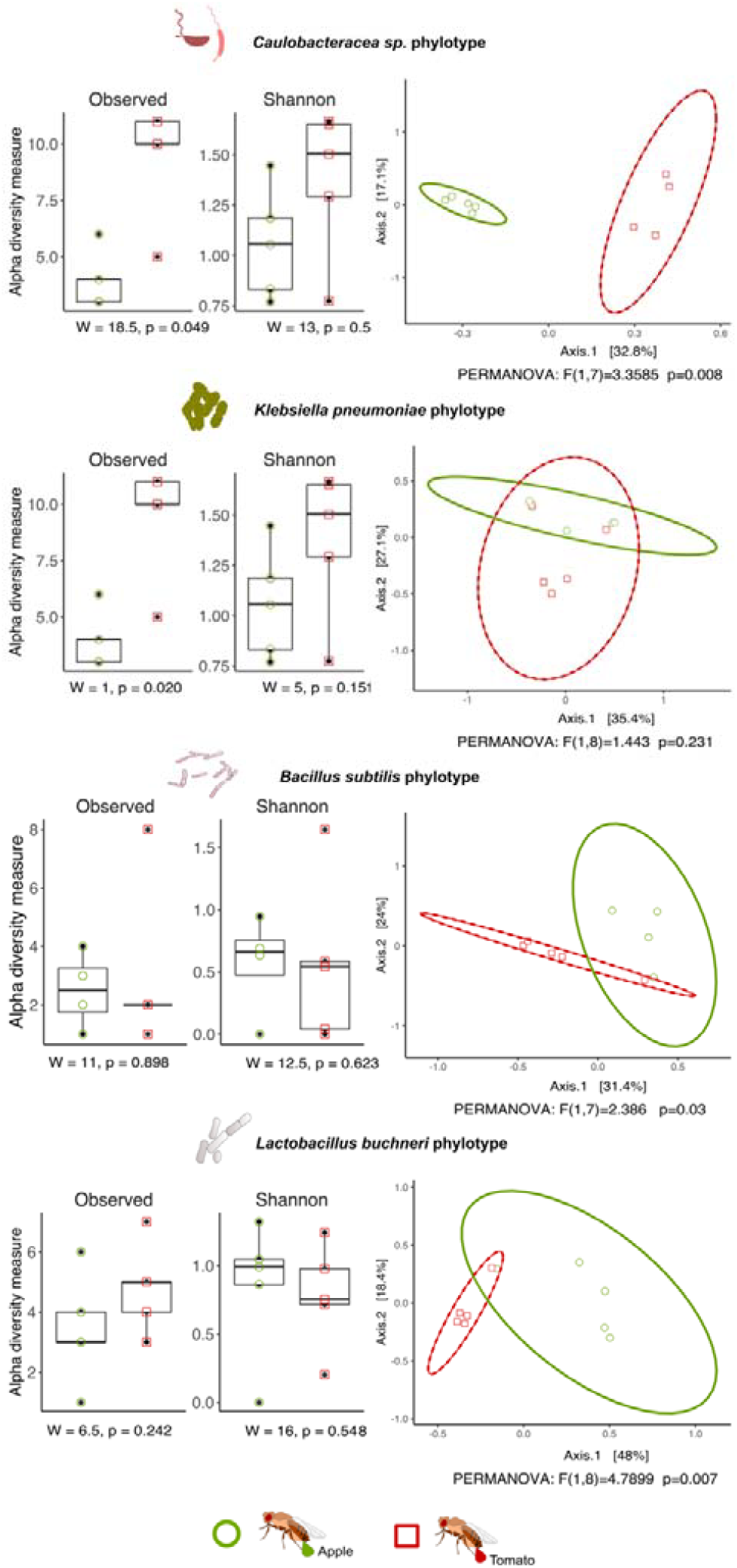
Microdiversity in bacterial symbionts of *Drosophila melanogaster* differs between breeding environments. Strain level diversity was analyzed in four selected bacterial phylotypes found in the fecal material of female *Drosophila melanogaster* reared in both apple and tomato environments. *Alpha*-diversity is shown as Observed and Shannon diversity, and environmental effects were analyzed by means of Wilcoxon rank sum test. Principal Coordinate Analysis (PCoA) based on Bray-Curtis dissimilarity was used to visualize differences in strain community composition (*beta-diversity*), ellipses represent 95% confidence. PERMANOVA were applied to identify influences of the insect breeding environment.

### Symbiont community composition changes during *Drosophila* development under *autochSI* and *allochSI* conditions

Next, we investigated the consequences of autochthonous (*autoch*SI) versus allochthonous symbiont inoculation (*alloch*SI) for microbiota composition in the larval environment. While *autoch*SI conditions in a tomato environment resulted in substrate bacterial and fungal microbiota similar to the initial fecal symbionts, *alloch*SI in an apple environment leads to a quite distinct microbial community (Fig 5). The shift in symbiont composition during larval development on the apple substrate was characterized by an increase in the relative abundance of *Pichia* yeasts (Fig 5) and the dominance of the Caulobacteraceae phylotype (Fig 5).

**Fig 5.**
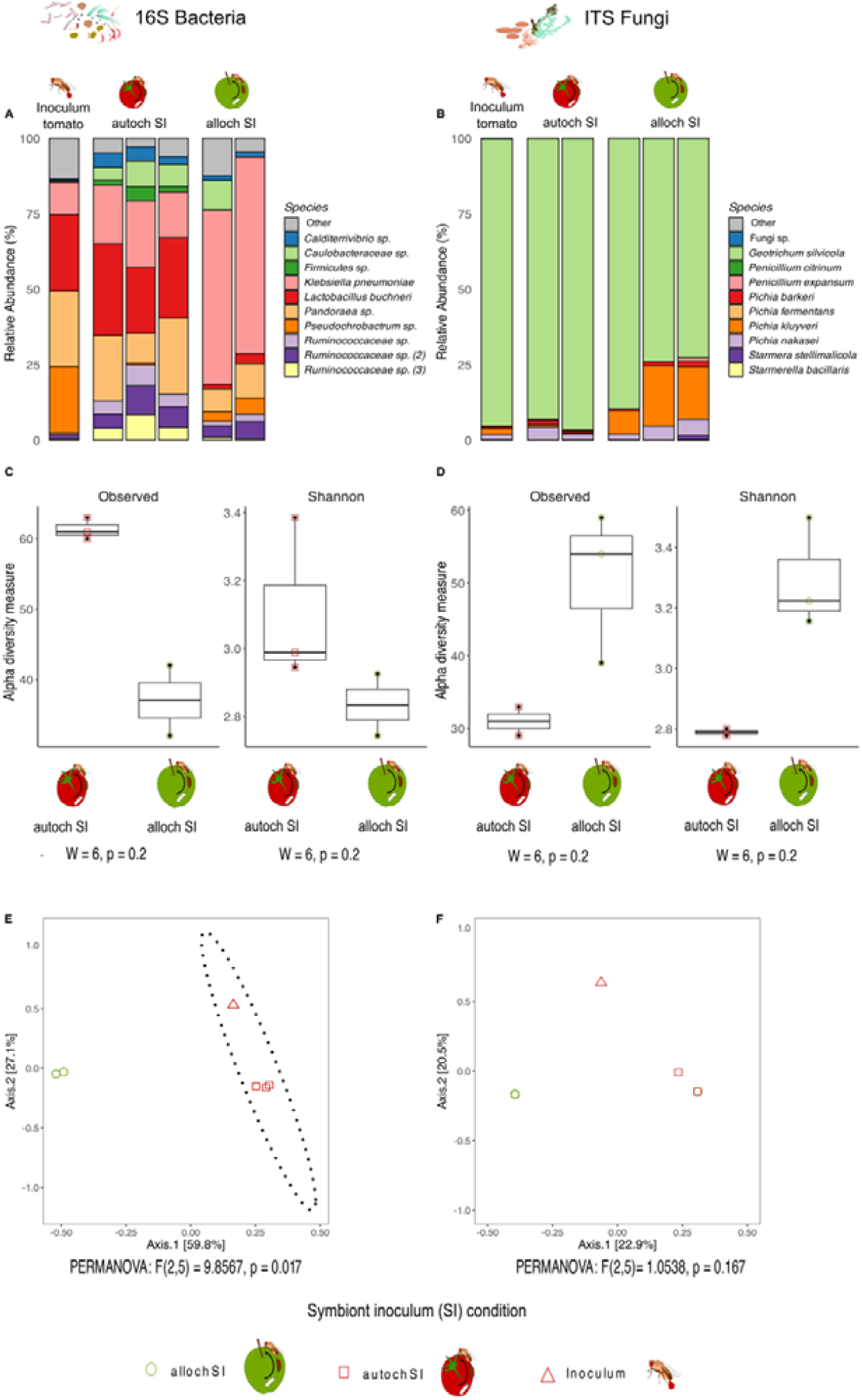
Allochthonous microbiota transmission alters larval *Drosophila melanogaster* symbiont composition. Relative abundance of the 10 most abundant bacterial (A) and fungal (B) symbionts in the fecal inoculum (single stacked bars, left) and on the autochthonous (tomato) and allochthonous (apple) substrates following larval development. *alpha*-diversity (Observed and Shannon) of symbionts following autochthonous and allochthonous inoculum and larval development (C, D). Principal Coordinate Analysis (PCoA) based on Bray-Curtis dissimilarity was used to visualize differences in bacterial and fungal community composition (*beta*-diversity) following larval development on the autochthonous (tomato) and allochthonous (apple) substrates, ellipse represent 95% confidence. PERMANOVA was applied to identify influences of the breeding substrate (E, F).

### Source-specific contribution of essential amino acids (EAA) during *Drosophila* development depends on symbiont inoculum conditions

We performed compound specific isotope analysis of EAA in two contexts: 1) the breeding substrate after fly eclosion and 2) freshly emerged *Drosophila* females. This analysis aimed to characterize changes in EAA composition within the habitat during larval development and to identify the sources of these EAA – whether derived from bacteria, fungi, or plant substrates (apple or tomato) – utilized by or available to the insects.

During *Drosophila* larval development, the EAA profiles of both plant substrates shifted toward profiles resembling those of bacteria and fungi (Fig 6A, Fig S4). However, compared to apple, tomato substrates tended to be more dominated by bacteria-derived EAA, particularly under *autoch*SI conditions. Notably, EAA profiles on both substrates were influenced by symbiont inoculum conditions (Fig 6A). On apple, *alloch*SI conditions caused a pronounced shift of substrate EAA profile toward bacteria derived EAA.

**Fig 6.**
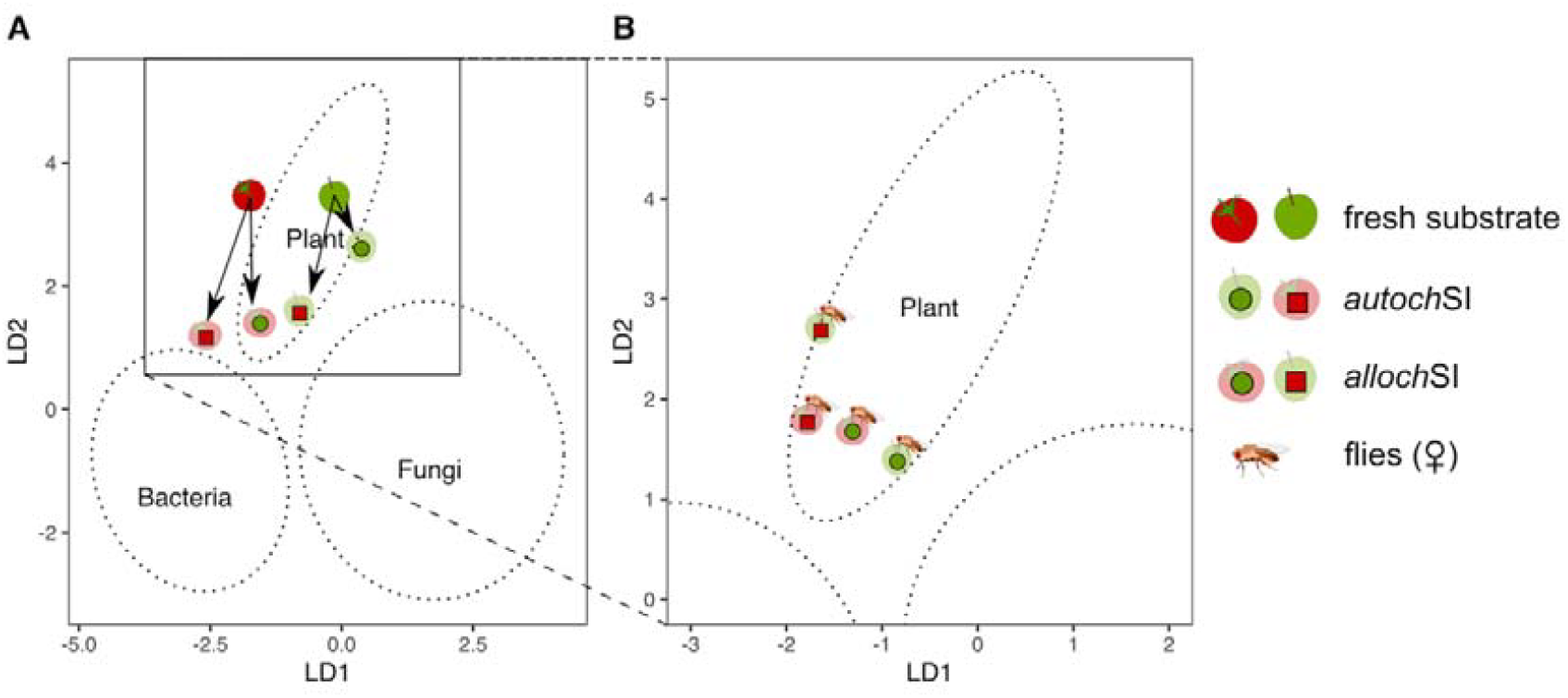
Symbiont inoculum conditions affect essential amino acid (EAA) flow during *Drosophila melanogaster* development. (A) Centroids calculated based on a linear discriminant analysis (LDA) of EAA-specific δ^13^ C values of fresh tomato and apple, and substrates following larval development under different symbiont inoculum conditions. (B) Fly EAA profiles following development under autochthonous and allochthonous symbiont inoculum condition. The ellipses represent the positioning of EAA profiles from different plant, bacterial, and fungal sources, respectively. They were used as classifiers in LDA (64). See Figure S4 for polygons depicting variation in EAA profiles of substrates and flies.

Interestingly, the EAA profiles of the flies showed only little resemblance to those of the substrates following fly eclosion; in fact, they remained similar to the EAA profiles of fresh substrate (Fig 6). Under allochSI conditions on tomato, fly and substrate EAA profiles following fly eclosion clustered similarly relative to each other. In contrast, fly EAA profiles under allochSI conditions on apple, resembled those of fresh tomato substrate. Notably, the EAA profiles of flies reared on a standard laboratory diet showed no overlap with those developed under semi-natural conditions on either apple or tomato substrates, irrespective of symbiont inoculum (Fig S4).

Independent of symbiont inoculum conditions, tomato substrate following fly eclosion and respective fly EAA profiles were lower on LD2 compared to fresh substrates, i.e. closer to microbial resources (Fig 6). The EAA profiles of both the tomato substrate and the eclosed flies tended toward a bacterial profile under autochSI (LD1, negative) and toward a fungal profile under allochSI conditions (LD1, positive). Apple substrate and the respective flies did not show these similarities in EAA profiles. Compared to fresh apple substrate EAA profiles of apple substrate after fly eclosion followed the same vector as the tomato substrates post eclosion. EAA profiles of flies that developed on apple did not follow these vectors.

Regardless of the plant substrate and symbiont inoculum conditions, both tomato and apple consistently served as the primary source of EAA for flies, contributing on average between over 50% to more than 75% (Fig 7). When developing in an apple environment, bacteria contributed slightly more than fungi to the EAA in flies that fed and grew as larvae under both autochSI and allochSI conditions. However, under allochSI conditions, the relative contribution of bacterial and fungal EAA dropped to well below 20%, resulting in the contribution of the apple substrate increasing to more than 75%. In the tomato environment, symbiont inoculum conditions hardly affected EAA incorporation by the larvae.

**Fig 7.**
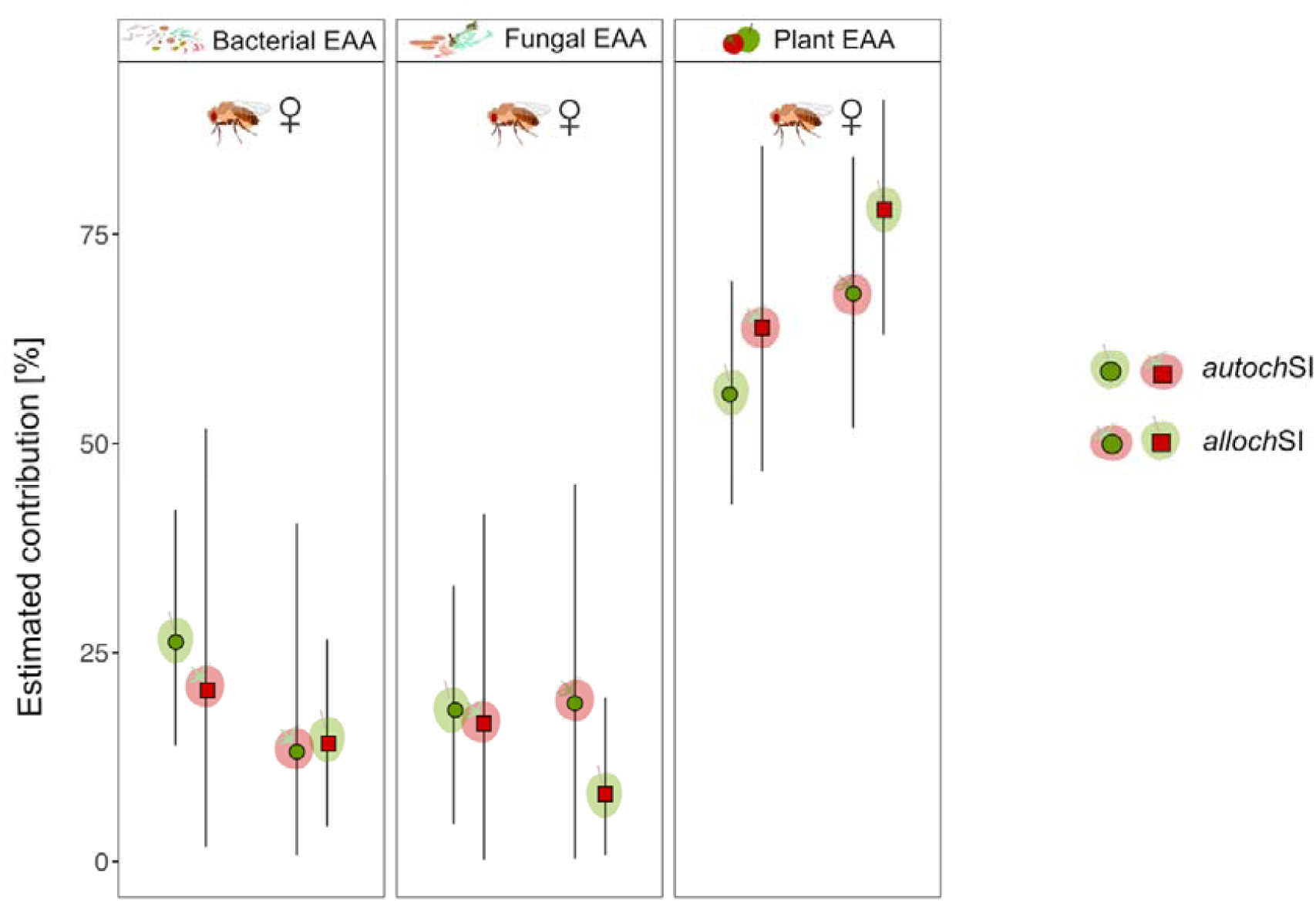
Symbiont inoculum conditions influence essential amino acid (EAA) uptake in *Drosophila melanogaster* larvae. Estimated relative contribution (%) of bacteria, fungi, and the plant substrates to EAA assimilated by larvae under autochthonous and allochthonous symbiont inoculum conditions. EAA were extracted from freshly eclosed female flies. Estimates are based on a Bayesian mixing model of mean-centered δ^13^ C values and displayed with 95% confidence interval.

### Expression of *Drosophila* life-history traits depends on symbiont inoculum conditions

We investigated whether environment-dependent differences in fecal symbionts and observed shifts in microbiota composition in response to allochSI impact Drosophila larval development. In a series of independent experiments (see Methods for details), we found a consistent pattern of effects of autochSI versus allochSI conditions on adult fly body weight and development time, respectively. In the presence of symbionts from an apple environment, fly weight remained consistent regardless of sex or symbiont inoculum conditions (Fig 8). Interestingly, larval development was faster under allochSI conditions compared to autochSI conditions. However, in an apple environment, allochSI conditions resulted in a significant decrease in body weight and simultaneously prolonged development times in both female and male flies (Fig 9). Given the significant effect of the statistical interaction between breeding substrate and symbiont origin on both body weight and development time (Table 1 and 2), these results indicate a symbiont-environment interaction.

**Table 1.** Type III ANOVA result for a general linear model testing predictors of z-transformed adult *Drosophila melanogaster* weight (mg).

| Explanatory variable | Sum of squares | DF | F-value | p-value |
| --- | --- | --- | --- | --- |
| Substrate | 5.82 | 1 | 9.08 | 0.003 |
| SI condition | 73.89 | 1 | 115.29 | <0.001 |
| Sex | 88.28 | 1 | 137.74 | <0.001 |
| Substrate x SI condition | 50.06 | 1 | 78.11 | <0.001 |
| Substrate x Sex | 2.89 | 1 | 4.52 | 0.034 |
| SI condition x Sex | 13.25 | 1 | 20.67 | <0.001 |
| Substrate x SI condition x Sex | 6.87 | 1 | 10.72 | 0.001 |
| Residuals | 505.03 | 788 |  |  |

**Substrate:** apple or tomato. **SI:** symbiont inoculum condition, either autochthonous or allochthonous

**Table 2:** Type III ANOVA result for a linear mixed model testing predictors of Box-Cox transformed development time in *Drosophila melanogaster* (days). Experimental run was specified as random factor.

| Explanatory variable | Sum of squares | DF (num) | DF (den) | F-value | p-value |
| --- | --- | --- | --- | --- | --- |
| **Substrate** | 8.547e-05 | 1 | 806.083 | 411.179 | <0.001 |
| **SI condition** | 9.017e-06 | 1 | 806.025 | 43.380 | <0.001 |
| **Sex** | 4.530e-06 | 1 | 806.022 | 21.795 | <0.001 |
| **Substrate x SI condition** | 1.829e-05 | 1 | 806.086 | 87.994 | <0.001 |
| **Substrate x Sex** | 4.901e-08 | 1 | 806.036 | 0.236 | 0.627 |
| **SI condition x Sex** | 9.406e-07 | 1 | 806.013 | 4.525 | 0.034 |
| **Substrate x SI condition x Sex** | 9.983e-07 | 1 | 806.126 | 4.803 | 0.029 |

**Substrate:** apple or tomato. **SI:** symbiont inoculum condition, either autochthonous or allochthonous

**Fig 8.**
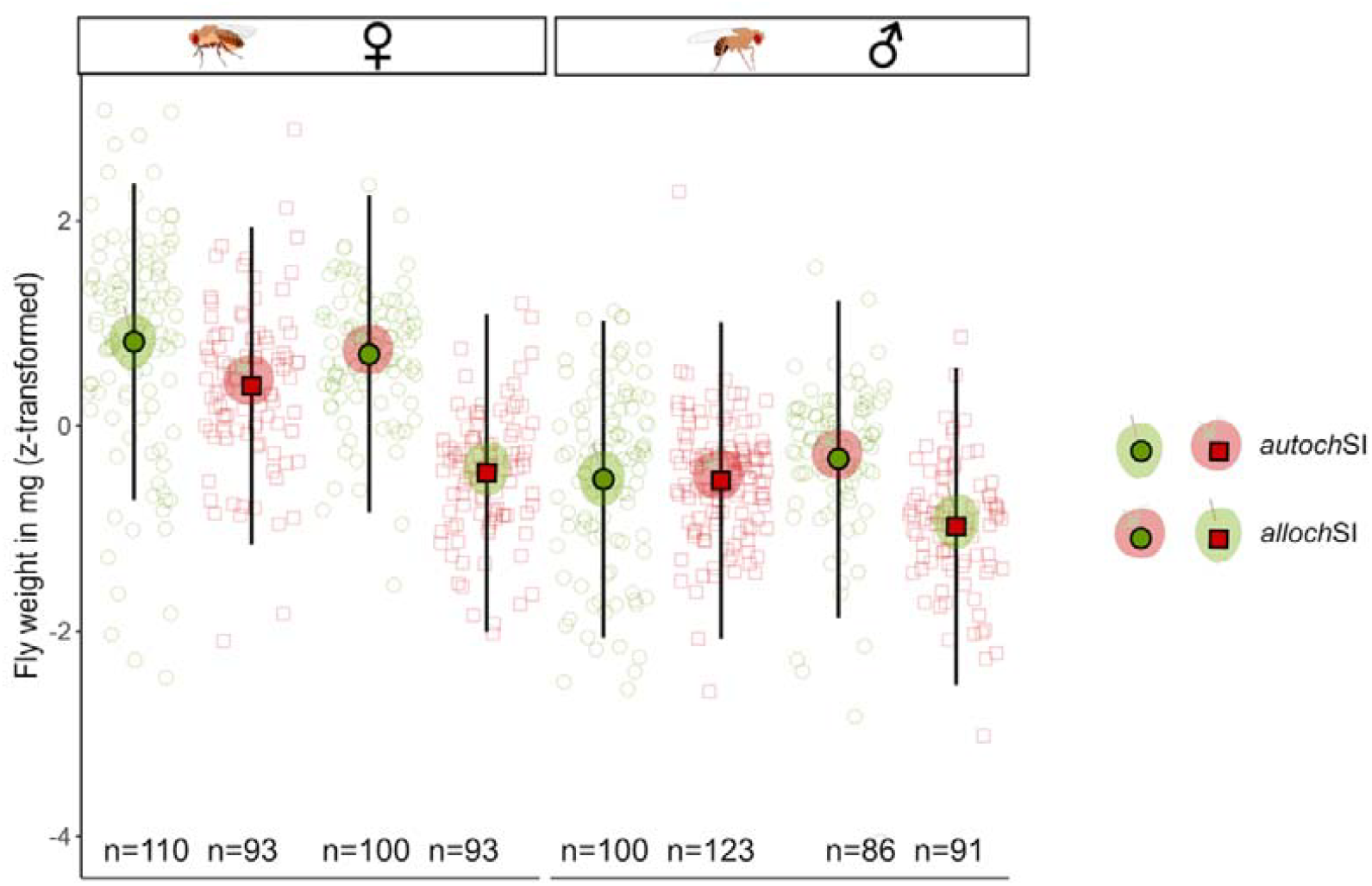
Biomass gain during development of *Drosophila melanogaster* larvae is affected by symbiont inoculum conditions. Body weight of freshly emerged male and female *Drosophila melanogaster* reared under different symbiont inoculum conditions. Larvae developed on either apple or tomato substrates inoculated with fly-derived fecal microbiota from an autochthonous (*autoch*SI) or allochthonous (*alloch*SI) environment. Raw data were z-transformed to account for baseline differences in average body weight across four experimental runs. Estimates (filled symbols) and corresponding 95% confidence intervals were generated using a general linear model. *n* indicates the number of individual flies measured. See Table 1 for Type III ANOVA.

**Fig 9.**
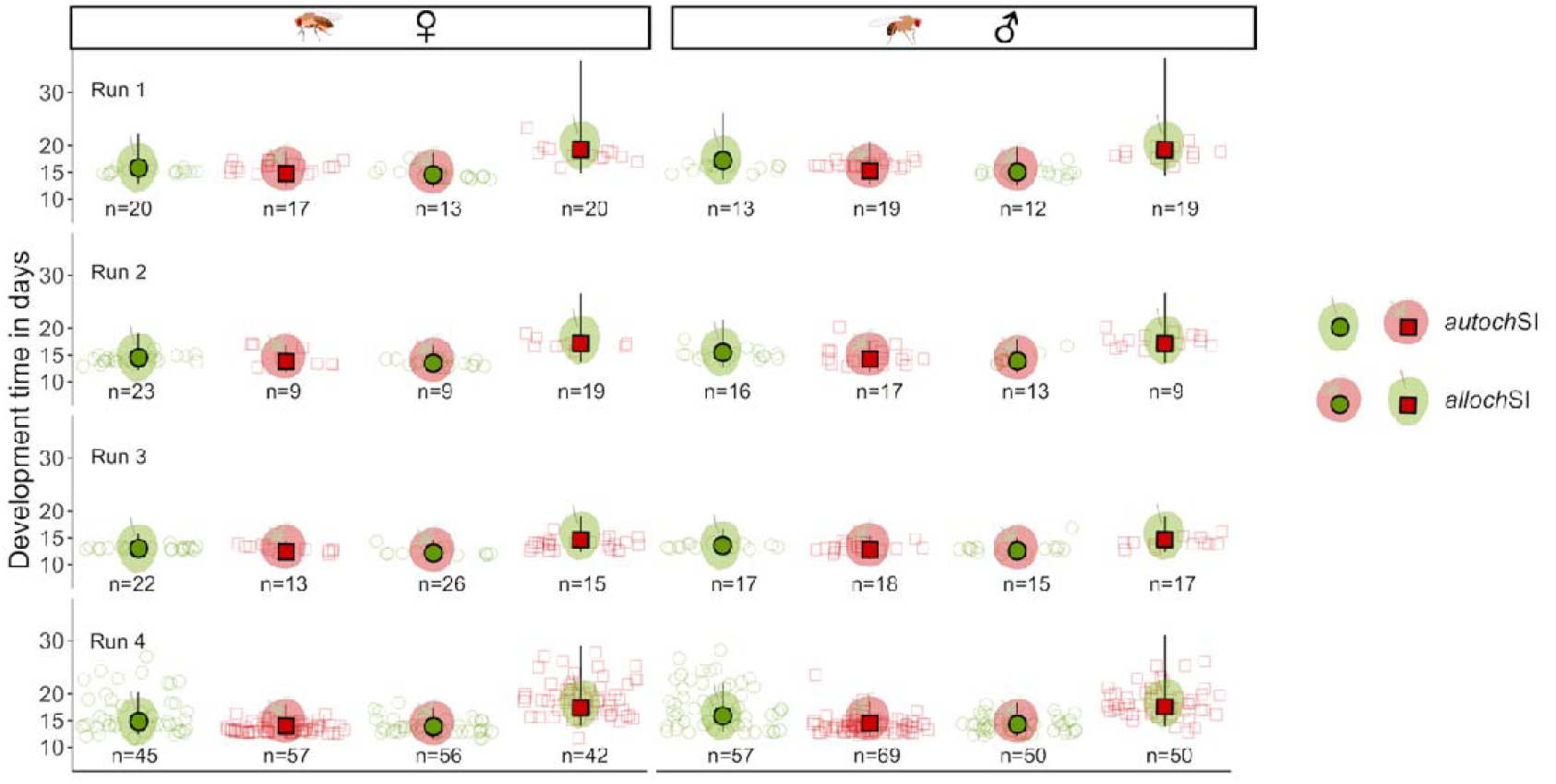
Development time of *Drosophila melanogaster* larvae is affected by symbiont inoculum conditions. Development time of male and female *Drosophila melanogaster* reared on apple and tomato substrates inoculated with fly fecal microbiota from either autochthonous (*autoch*SI) or allochthonous (*alloch*SI) environments. Development time was measured in days from first instar larva to adult eclosion. Estimates were generated for each combination of substrate, symbiont inoculum, and sex using a linear mixed-effects model that corrected for variation across experimental runs (*n* = 4 per condition, *n* = 2 sexes). Error bars represent 95% confidence intervals. *n* indicates the number of individual flies for which development could be recorded. See Table 2 for Type III ANOVA.

The relatively long development time and low body weight of female flies reared as larvae under *alloch*SI conditions on apple correlate with the highest contribution of plant-derived EAA and the lowest contribution of fungus-derived EAA to their EAA profile (Fig 7, Fig S5). Variation in bacteria-derived EAA provides limited explanatory power for differences in insect life-history traits.

## Discussion

In this study, we provide evidence of symbiont-environment interactions affecting the developmental success of *D. melanogaster*. We define symbiont-environment interactions as a situation where the influence of microbial symbiont composition on host development depends on the ecological context of the breeding environment. We explored this experimentally by simulating autochthonous and allochthonous symbiont scenarios within the lifecycle of *D. melanogaster* (Fig 1). To bridge the gap between laboratory and natural conditions, we established fly populations in semi-natural microcosms using fruit substrates (apple and tomato), where field-derived microbiota was transmitted and maintained across multiple generations. This approach allowed us to observe consistent environment-dependent variation in symbiont communities, both in adult females and in larval developmental substrates. By linking symbiont composition, essential amino acid uptake, and life-history trait expression, we present a novel approach to investigate environmental heterogeneity shaping host-microbe interactions in the *D. melanogaster* model system.

### The breeding environment determines bacterial and fungal symbiont diversity of *D. melanogaster*

Our results experimentally verify the finding of observational studies (28,32,69) that the plant substrate defining the *Drosophila* breeding habitat is a fundamental driver of the microbiota forming a symbiotic relationship with both the adult flies and the larvae. The deposition of fecal material near oviposition sites is a primary route through which larvae acquire microbial symbionts (25,27). The fecal material deposited by female flies reared in a tomato environment tended to contain more diverse bacterial communities than those reared in an apple environment. Despite variation in bacterial *alpha*-diversity across years and transmission histories (Fig S3), substrate environment consistently explained differences in community composition (*beta*-diversity). First, this demonstrates a robust, environment-dependent association of bacterial symbionts across generations. Second, it challenges the common view of *Drosophila* microbiota as “simple” e.g., Ludington and Ja (70), revealing substantial variation shaped by ecological context.

While several Enterobacteriaceae phylotypes and a Caulobacteraceae phylotype dominated the fecal microbiota of flies from an apple environment, the community composition in the fecal material of flies from a tomato environment appeared more even with higher proportions of Burkholderiaceae and Lactobacillaceae, among others. Except for the Caulobacteraceae phylotype, the families and phylotypes identified in both populations generally align with those reported to be fly associated in nature by other studies (14,18,71– 74). Surprisingly, we did not detect a dominance of Acetobacteraceae, often reported in other studies (18,71,72,74). Acetobacteraceae phylotypes have previously been shown to establish a stable association with *Drosophila*, supported by gut-specific conditions that promote their growth (24). Some of the mechanisms that enable such stable gut-bacterial associations in *Drosophila* have only recently been revealed (75). These mechanisms involve a complex interplay of host–microbe and microbe–microbe interactions that shape a specialized niche in the proventriculus of adult flies, ultimately favoring the persistence of microbial taxa capable of propagating and establishing stable associations with the host. Dodge et al (75) links the establishment of stable symbiosis to chemical alterations of the fly gut by microbial partners. As our study demonstrates, the pool from which microbes may be selected varies greatly across environments, leading to the establishment of distinct symbiont communities. Accordingly, we expect that the mechanisms identified by Dodge et al. (75)are sufficiently adaptable to respond to the substrate environment’s influence on shaping the microbiota ingested by flies. The dominant bacterial taxa transmitted by flies in our study likely possess traits that enable their selection through these mechanisms, possibly facilitating their integration into stable, habitat-dependent mutualistic relationships (see also below).

In addition to environment-dependent differences in phylotype diversity, a more detailed analysis of amplicon sequence variants (ASVs) revealed microdiversity within several bacterial phylotypes, including *Klebsiella pneumoniae, Bacillus subtilis, Lactobacillus buchneri* and a Caulobacteraceae phylotype. Notably, both number of strains and strain composition analyses showed, that the composition of ASVs, within single phylotypes in the fecal material of female *Drosophila*, can vary depending on the substrate environment in which the flies were kept. In a study on the honeybee gut microbiome, Baud et al. (76) link microdiversity differences between nursing and foraging bees to dietary shifts, specifically the cessation of pollen consumption in foragers. Similarly, a cautious interpretation of our results could be that within-phylotype genetic variation in a given environment undergoes selection, favoring variants that are best adapted to thrive in and be transmitted by flies between substrate patches of a specific kind. This finding thus highlights two important novel aspects: First, microdiversity in bacterial symbionts must be considered in *Drosophila* microbiome analyses to uncover their potential impact on host phenotypic trait expression. Second, the ecological context – specifically, the breeding substrate environment – should be regarded as a potential selective filter for ASV composition. The extent to which host control mechanisms (see above) additionally contribute to the maintenance and differentiation of microdiversity remains to be determined.

Fungal symbiont diversity in the feces of female *Drosophila* was also influenced by the substrate environment. The *alpha*-diversity analysis revealed greater fungal diversity in the fecal material of flies reared in an apple environment compared to those in a tomato environment, while *beta-*diversity analysis showed only marginal differences. Fungal symbionts in “apple” flies were dominated by frequently found species of *Starmerella* and *Pichia* (e.g.,23,25,76) which were nearly absent in the “tomato” flies. Most strikingly, *Geotrichum silvicola* overwhelmingly dominated the fungal community in the feces of flies from the tomato environments. *G. silvicola* was originally isolated from insects, including *Drosophila* (78). Furthermore, various *Geotrichum* species, including those that are plant pathogens, are frequently found in association with *Drosophila* (73,77,79,80). Since fly populations thrived in our tomato microcosms, *Geotrichum*, which exhibits morphological traits intermediate between yeasts and filamentous mold fungi (81), does not seem to be pathogenic for the insects.

Although fungi can persist in the flies’ gut system for an extended period (25), there is currently no evidence that they establish a long-term association without ongoing replenishment. This suggests that fungal symbiont communities are likely to change when the flies’ breeding environment changes. Supporting evidence comes from Gurung et al. (82), who found that fungi associated with *D. suzukii* were similarly associated to their breeding substrate, unlike their bacterial symbionts. Since we observed effects of the breeding substrate on both fly-associated bacteria and fungi, we cannot confirm that bacteria are more persistently associated with *D. melanogaster*. A possible explanation could be that apples and tomatoes differ more significantly from each other in their properties (see below) than the different types of berries used by Gurung et al. (82). The plant substrate can be a very strong filter for the symbionts that establish in the flies’ breeding site, which becomes apparent when comparing symbiont community development under autochthonous inoculation (*autoch*SI) conditions to that under allochthonous inoculation (*alloch*SI) conditions (Fig 1), as discussed below.

Under *alloch*SI conditions, we observed a significant shift in bacterial symbionts from “tomato” flies that established on apple substrate. Unlike the *autoch*SI conditions, which led to a community similar to the initial inoculate, bacterial diversity on apple was notably reduced and characterized by a pronounced shift towards a Caulobacteraceae phylotype. In contrast, fungal phylotypes slightly increased under *alloch*SI conditions, however this was just a trend. This demonstrates how common environmental variation in the *Drosophila* habitat can lead to substantial changes in symbiont composition.

In conclusion, the breeding substrate environment of *Drosophila* influences both bacterial and fungal symbionts. This is evident for symbionts transmitted by adult flies from different habitats, as well as for those that grow in the presence of larvae under *alloch*SI versus *autoch*SI conditions. While we can currently only speculate on the mechanisms underlying these differences, it likely involves significant variation in substrate acidity, water availability, and nutrient content. For example, we found the substrate pH in apple and tomato substrates to vary between 2.6 - 3.2, and 3.6 - 4.4, respectively (S3 Table). Oakeshott (31) identified a correlation between substrate-dependent differences in pH, the abundance of bacteria and different kinds of yeast fungi, and the distribution of various *Drosophila* species.

Although substrate properties directly shape microbe–microbe interactions that influence host performance(22,83–86), both adult and larval flies can further modulate symbiont composition (e.g. Stamps et al. 2012). This has recently been shown in *Ceratitis capitata* (Tephritidae) harboring a microbiota strongly influenced by host fruit but regulated through life-stages and host functional needs (87). Such effects may arise through individual immune traits (88), the gut’s capacity to lyse or promote specific microbes (24,75,89), and cooperative interactions within larval aggregations (44,90,91). It remains unknown if host traits and system properties allow *D. melanogaster* to exert a density-dependent “leash” on its microbiome (9).

### Essential amino acid contribution to *D. melanogaster* tissue depends on plant substrate-symbiont combination

One of the most relevant “services” of symbionts is to promote host fitness through improving dietary conditions, such as biopolymer degradation, provisioning of essential vitamins and amino acids (2,92). A similar importance of symbiont microbes providing essential amino acids (EAA) in an otherwise (artificial) low-protein environment has been demonstrated for both larval *Drosophila* development, and adult lifespan and reproduction (19,93). To the best of our knowledge, combining our semi-natural experimental approach with stable isotope analysis of EAA in freshly emerged flies has not been done so far. Thus, we were able to reveal for the first time the actual sources of EAA incorporated during larval development. High proportion of EAA found in the flies indeed came from the plant substrate. This proportion even increased under *alloch*SI conditions, whereas the proportion of EAA contribution from the plant to the insect tissue was always higher in the presence of symbionts from a tomato environment. Possibly, the environmental context-dependent shifts in EAA contributions from the three sources, plant, bacteria and fungi, is due to substrate-induced changes in the prevalence of nutrient-rich, nutrient-poor, unpalatable, or even pathogenic symbionts.

Interestingly, EAA profiles of flies did not resemble the EAA profile of the substrate after larval development (see Fig 6). Again, differences in symbiont communities may create nutritional niches that constrain EAA uptake. Furthermore, larvae may directly influence EAA uptake through selective feeding (94). While the EAA profile of the substrate reflects the endpoint of microbial succession during larval development, the larval EAA profile results from the dietary intake of EAA throughout ontogeny. Therefore, the substrate EAA profiles cannot represent microbial dynamics during larval development. These, still unknown, dynamics in symbiont community succession – relative abundances and absolute population sizes – may additionally explain the mismatch between substrate and insect EAA profiles.

Lastly, we note that the EAA profiles of flies reared on an artificial laboratory diet showed no overlap with those of flies developing in natural breeding substrates, regardless of their associated symbionts. Although we can, beyond variation in body size (see below), only speculate on the consequences for *Drosophila* nutritional physiology (95), our findings show that the quality of microbial partners depends strongly on the breeding substrate. This makes explicitly including ecological context essential for understanding the selective environments that shape the evolution of both *Drosophila* life-history (21) and its mutualisms with microorganisms (see below).

### *D. melanogaster* phenotypic trait expression is shaped by symbiont-environment interactions

Symbiont-environment interactions, shaped by variation in *D. melanogaster* microbiota *across* habitats and substrate-dependent dynamics *within* substrate patches, manifest in significant differences in two key life-history traits: larval development time until adult eclosion and adult body weight. Fly body weight was not uniformly affected by *alloch*SI conditions, instead, our experiment revealed that larval weight gain was relatively lower when symbionts originated from a tomato environment. In contrast, the effect of symbionts from “apple” flies did not vary significantly between *autoch*SI and *alloch*SI conditions. Development time was prolonged under *alloch*SI conditions in the presence of microbiota from “tomato” flies. Interestingly, when larvae were reared with symbionts from “apple” flies under *alloch*SI conditions, their development time was even accelerated. These findings suggest that when substrate availability varies and female flies transmit allochthonous microbes, symbiont-environment interactions can lead to both fitness gains – such as faster development without a reduction in body mass – and fitness losses, as seen in slower development and reduced body mass under *alloch*SI conditions. Therefore, symbiont-environment interactions offer a new conceptual framework for better understanding microbiota-driven phenotypic variation in *Drosophila*, which in consequence determines the width of the reproductive niche of the entire symbiotum.

As discussed above, the most striking outcome of the symbiont-environment interaction observed in this study is the massively reduced body size and prolonged development time under *alloch*SI conditions on apple. Possibly, this could be attributed to the higher intake of EAA from the plant substrate, relatively to the intake from the microbial symbionts. Since yeast fungi alone have been shown to provide a sufficient nutrient source that promotes larval growth (25,94), the stability in body weight observed with symbionts from apples may be due to a largely unaltered EAA contribution from fungi, even under *alloch*SI conditions. The striking dominance of plant-derived EAA in flies that developed under *alloch*SI conditions on apple may negatively impact larval development through several potential mechanisms. One possibility is that symbiont-derived EAA and other essential nutrients, such as steroids, become scarce due to reduced microbial (fungal?) growth in the new environment. This forces larvae to rely on plant EAA, which may exist in lower concentrations and be harder to extract from the plant matrix. Additionally, certain *alloch*SI conditions may alter the symbiont community composition, promoting the proliferation of low-quality or even pathogenic microbes (22). This shift could trigger secondary metabolite-mediated dysbiosis, potentially disrupting larval development (96,97). The observed changes in larval weight gain over time could also result from a complex interplay between changes in symbiont communities and host behavior. For instance, microbe-induced changes in feeding behavior (50) and the varying prevalence of symbionts with beneficial or detrimental traits might collectively influence development outcomes, and thus the reproductive niche space of the host (98). To deepen our understanding of these dynamics, future research should focus on isolating key microbial taxa, assessing their individual and combined effects on host traits, and applying synthetic communities under controlled yet ecologically relevant conditions that mirror the environments in which host-microbe interactions naturally evolve (15).

### Implications for understanding the evolution of *Drosophila*-microbe interactions

The substrate-dependent differences in microbial community composition across breeding environments indicate that mutualistic dependence between *Drosophila* and its microbiota is largely facultative. Although larval development and adult reproduction remain viable across a range of microbial assemblages (99), not all symbiont communities contribute equally to host development. Variation in fitness outcomes under *alloch*SI conditions suggests that microbial partners differ in the relative benefit depending on ecological context. This supports the view that *Drosophila* interacts with its microbial partners along a continuum of mutualistic quality, with weak dependence on individual phylotypes likely limiting the evolution of specialization (10).

Crucially, while *Drosophila* larvae require microbial input for successful development, this dependency is functional rather than obligate in the evolutionary sense. Development relies on microbial functions, such as essential amino acid provisioning, which can be fulfilled by a wide range of bacteria and fungi. This positions *Drosophila* as an obligate generalist that cannot develop without microbial partners, but partner breadth is potentially broad (10,11). For microbial symbionts, the unpredictability of fly breeding environments and thus the lack of consistent transmission between host generations limit the evolution of stable associations and cooperative traits. Most identified symbionts can associate with a variety of invertebrate hosts and persist independently in the environment e.g., Carramachi et al. (100). Altogether, these findings suggest that ecological context constrains the evolution of specialization in *Drosophila*-microbe interactions and, through broad partner breadth, stabilizes a functionally essential yet evolutionarily loose form of mutualism.

Despite limited specialization in *Drosophila*-microbe interactions, a deeper understanding of symbiont-environment dynamics offers a framework for identifying microbial taxa with the potential to form stable, long-term associations across diverse ecological contexts. For example, a single variant of the *Lactobacillus buchneri* phylotype was detected in the fecal material of flies from diverse microcosm environments beyond apple and tomato, including plum, grape, lemon, and onion (Riedel & Rohlfs 2025 *unpublished*). The broad occurrence of this variant suggests it may have the potential to evolve into an obligate specialist under specific ecological conditions, a hypothesis that could be tested experimentally by evaluating its functional contributions to larval development across diverse substrates. A similar case was reported for *Acetobacter thailandicus* in *D. melanogaster* (24), where an isolate of *A. thailandicus* colonized the gut of several *D. melanogaster* genotypes, proliferating and providing sufficient nutrients to sustain the host on otherwise sterile figs. However, this interaction was only tested within the fig environment. Alternatively, rather than relying on specialized microbial partnerships, *Drosophila* may benefit from actively shifting breeding substrates to replace symbionts with limited functionality (10,11). As outlined previously, while *Drosophila* can influence its microbiota through individual behavior, social interactions, and physiology (18), it remains unclear to what extent this control can override the strong filtering imposed by the breeding environment. To disentangle relative roles of host control and environmental filtering will be essential for linking natural ecological dynamics to the evolution of host-microbe associations in this model system.

Lastly, our findings raise the possibility that variation in symbiont community composition contributes to host adaptation in novel environments (43,98,101). The faster development of flies under *alloch*SI conditions on tomato, compared to those reared with autochthonous microbes, suggests that non-local microbial communities may sometimes confer unexpected nutritional or physiological advantages. These results point to a broader role for flexible generalist symbioses in promoting adaptive plasticity, potentially explaining how *Drosophila* populations maintain breeding niche breadth across diverse and ephemeral plant substrates and may even expand into previously unexploited resources (102).

Our study highlights how ecological context shapes nutritional interactions between *D. melanogaster* and its microbial associates. Symbiont-environment interactions reveal context-dependent costs and benefits of symbiont-dependent development, underscoring the contingencies of generalist associations. This framework offers new directions for exploring the ecological limits of evolutionary processes in host-microbe interactions.

## Acknowledgments

We thank Stefan Scheu for sharing knowledge and lab resources. We thank Guido Humpert and Nico Binnemann for technical support, and Jens Dyckmans for compound-specific isotope measurements of amino acids. Data availability of this study was supported by the Data Science Center of the University of Bremen (DSC@UB) funded by the State of Bremen.

## Data availability statement

All data associated with this study were brokered through the German Federation for the Curation of Biological Data (GFBio). The raw amplicon sequencing data for this study have been deposited in the European Nucleotide Archive (ENA) at EMBL-EBI under accession number PRJEB96936 and PRJEB96953. Biological trait data (development time, weight), amino acid δ13C isotopic profiles, and modeled nutrient sourcing outputs are available form PANGAEA, https://doi.org/10.1594/PANGAEA.989940^103^.

