## Supporting_Information for "Ecological context shapes microbial contributions to nutrition and development in *Drosophila melanogaster*"

### S1 Text. Media used for fly rearing and exemplary plating

Media composition of a) laboratory diet for Drosophila melanogaster (artificial medium), b) selective growth medium for bacteria, Luria Broth (LB) and c) selective growth medium for fungi, Yeast Glucose Chloramphenicol (YGC).

Media recipes

a) Laboratory diet recipe:
- 12 g agar (Agar-Agar, Kobe I, powdered, Carl Roth GmbH + Co. KG)
- 50 g refined sugar (Diadem, Aldi)
- 10 mL methyl-4-hydroxybenzoat (10% in ethanol) (Sigma Aldrich)
- 10 mL sorbic acid (10% [w/v] in ethanol) (Carl Roth GmbH + Co. KG)
- 250 g apple puree (Sweet Valley, Aldi)
- 50 g cornmeal (Bauckhof Mühle)
- 70 g brewers' yeast (Leiber GmbH)
- 600 mL water

b) Luria Broth agar (selective for Bacteria) recipe:
- 20 g LB broth (Carl Roth GmbH + Co. KG)
- 100 mg nystatin (Carl Roth GmbH + Co. KG)
- 1000 mL water (DI)
- 14 g agar (Agar-Agar, Kobe I, powdered, Carl Roth GmbH + Co. KG)

c) Yeast Extract Glucose Chloramphenicol agar (selective for fungi) recipe:
- 3 g yeast extract (Carl Roth GmbH + Co. KG)
- 3 g malt extract (Carl Roth GmbH + Co. KG)
- 5 g peptone (from soy) (Carl Roth GmbH + Co. KG)
- 10 g glucose (Fisher Chemicals)
- 1000 mL water (DI)
- 14 g agar (Agar-Agar, Kobe I, powdered, Carl Roth GmbH + Co. KG)
- 100 mg chloramphenicol (Carl Roth GmbH + Co. KG)

**Table S1. Microcosms used for fecal material sampling in five larval development experiments.** Each microcosm is defined by the collection year and the number of *Drosophila melanogaster* population transfers since establishment (see Materials and Methods). Bold numbers indicate replicates included in the developmental assay analyses after quality control; numbers in parentheses denote the initial number of experimental units prepared.

| **Symbiont inoculation (SI) conditions** | **Substrate type and replicates** | |
|---|---|---|
| | Apple | Tomato |
| March 2020 run • collection 2019, 5 transfers * | | |
| *autoch* SI | **38** (40) | **39** (40) |
| *alloch* SI | **34** (40) | **40** (40) |
| September 2020 run • collection 2019, 12 transfers * | | |
| *autoch* SI | **43** (50) | **36** (50) |
| *alloch* SI | **38** (50) | **32** (50) |
| March 2021 run • collection 2020, 5 transfers * | | |
| *autoch* SI | **44** (50) | **46** (50) |
| *alloch* SI | **44** (50) | **44** (50) |
| June 2021 run • collection 2020, 21 transfers † | | |
| *autoch* SI | 0 | **58** (60) |
| *alloch* SI | **58** (60) | 0 |
| October 2021 run • collection 2019, 26 transfers ‡ | | |
| *autoch* SI | **143** (150) | **142** (150) |
| *alloch* SI | **146** (150) | **126** (150) |

*autoch* : autochthonous, *alloch* : allochthonous, * Larval development only, † Substrate microbiota sampling only, ‡ Larval development and essential amino acid analysis

**Figure S1. Sufficient sampling depth visualized as rarefaction curves of fecal and substrate metabarcoding samples.**

Rarefaction curves of bacterial reads (A) and fungal reads (B) in fecal material samples from apple (green solid lines) and tomato (red dashed lines) fly populations. Rarefaction curves of bacterial reads (C) and fungal reads (D) in substrate samples, of apple (green solid lines) and tomato (red dashed lines) substrate, initially inoculated with tomato population fecal microbiota and sampled after fly development.

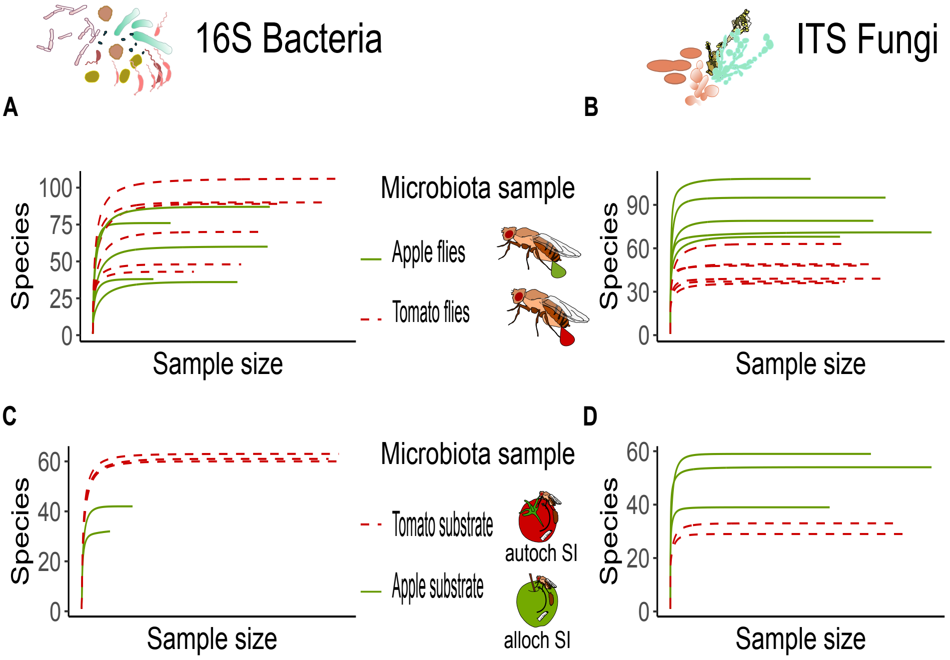

**Figure S2. Substrate environment determines Drosophila melanogaster symbionts.**

The most abundant Families (Bacteria) and Genera (Fungi) found in fecal material of flies from apple (green droplet) or tomato population (red droplet), over time (-1 first transfer event, -2 second transfer event etc.) and year of microcosm establishments (2019 and 2020 respectively) for bacterial (16S) and fungal (ITS) symbionts.; Percentages are given for all taxon groups over 1% of the relative abundance within sample.

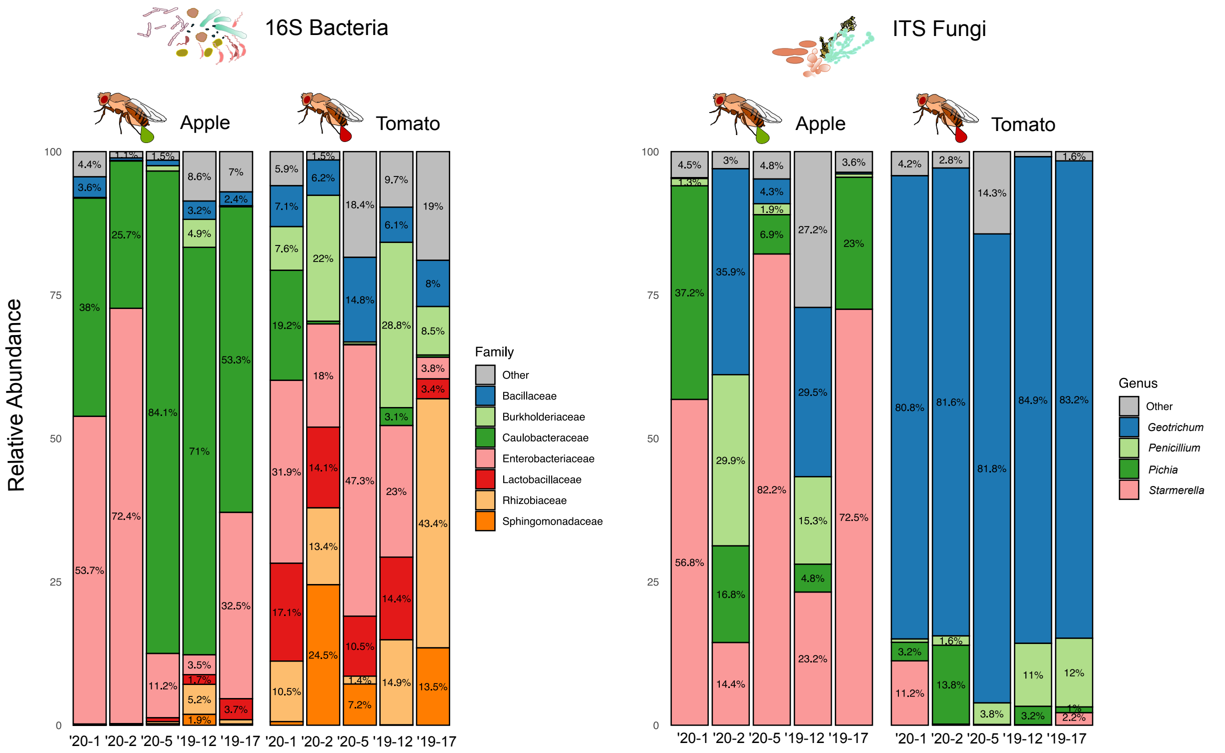

**Figure S3. Large differences in *Drosophila melanogaster* symbiont diversity among collection years and symbiont transfer events.**

Observed and Shannon diversity of symbiont communities in the fecal material of *Drosophila melanogaster* females from apple (green circles) or tomato (red squares) environments. The ITS2 region and the 16S ribosomal subunit were used as marker genes to characterize fungal and bacterial communities, respectively. The five sampling points are described with year of microcosm establishment ('20 or '19) and the number of transfers of the specific microcosm (-1, -2, -5, -12, -17) on the x-axis.

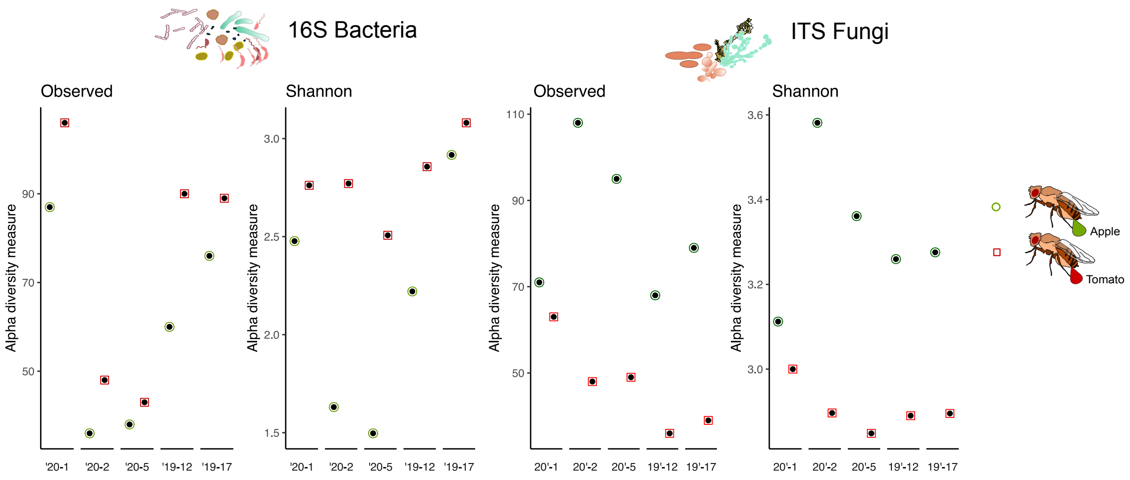

**Table S2. *Drosophila melanogaster* symbionts include phylotypes with multiple Amplicon Sequence Variants (ASV).** Over all 5 fecal samples, collected from 30 female "apple" or "tomato" flies, phylotypes are listed, that are represented by at least five ASV. Of these ASV only a subset is present in both "apple" and "tomato" fly microbiota; the number of strains unique to one fly environment type are listed.

| Phylotype name | Fecal microbiota of flies from apple environment | | Fecal microbiota of flies from tomato environment | |
|---|---|---|---|---|
| | No. strains | No. of which are unique | No. strains | No. of which are unique |
| Bacillaceae sp. | 6 | | | |
| Bacillus sp. | 6 | | | |
| *Bacillus subtilis* * | 7 | 4 | 9 | 6 |
| Bacteria sp. | 11 | 6 | 24 | 19 |
| Caulobacteraceae sp.* | 23 | 15 | 11 | 3 |
| Corynebacteriales sp. | 8 | | | |
| *Corynebacterium kutscheri* | 6 | | 6 | |
| *Klebsiella pneumoniae** | 8 | 0 | 22 | 14 |
| *Lactobacillus buchneri** | 10 | 5 | 10 | 5 |
| Pandoraea sp. | 12 | 5 | 13 | 6 |
| Proteobacteria sp. | 7 | | | |
| Pseudochrobactrum sp. | 6 | | | |
| Ruminococcaceae sp. | | | 12 | |
| Ruminococcaceae sp. (2) | 7 | 1 | 23 | 17 |
| Ruminococcaceae sp. (3) | | | 10 | |
| Virgibacillus sp. | 9 | 6 | 6 | 3 |

***)** Phylotypes selected for further analysis

**Figure S4. Substrate specific microbial growth changes AS composition of D. melanogaster developmental environment.**

Linear discriminant analysis of samples of flies (females, pools of 8), corresponding experimental units (developmental environment) and initial (fresh) substrate collected from larva development assay (October 2021, see Methods). Projected sample positions in the LDA space are connected with lines to form polygons. Flies raised on laboratory diet (Lab) and the respective substrate post-eclosion, had no added microbiota and are shown in green.

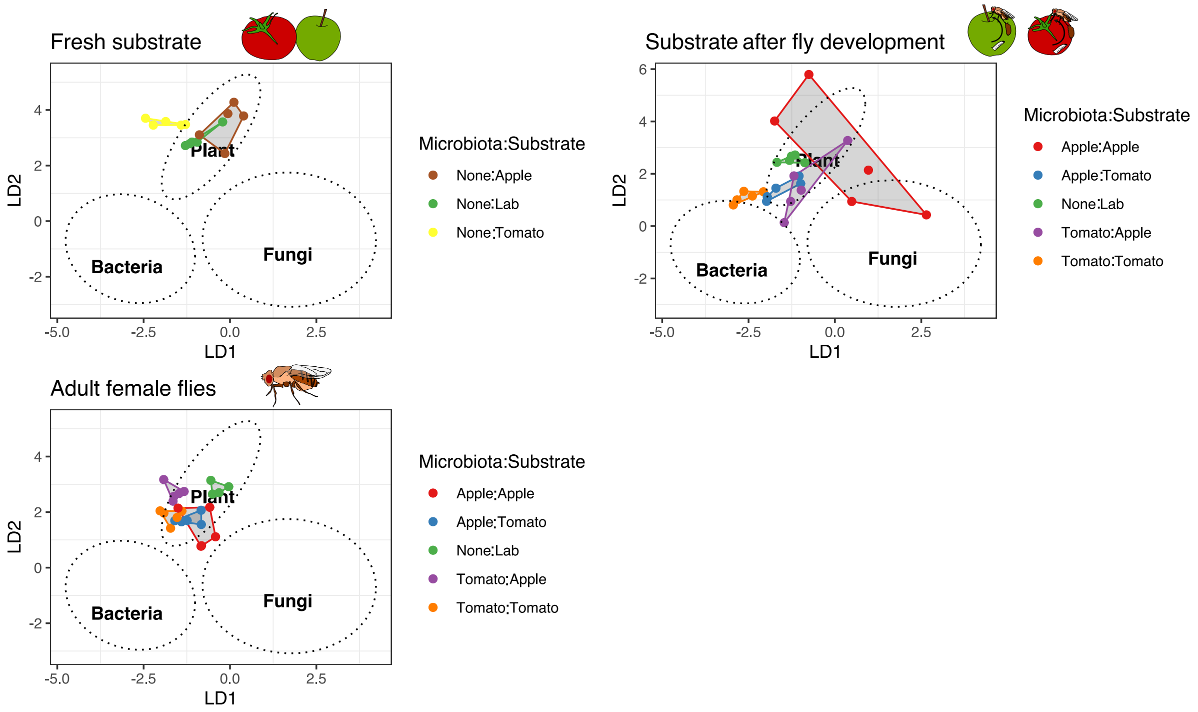

**Figure S5. Relative contributions of bacteria, fungus and plant-derived essential amino acids (EAA) in relation to female life-history traits in Drosophila melanogaster.**

Mean life-history trait values as modelled for run four of the larval development experiments (see Fig 8 and 9) by general linear model (z-transformed body weight) and generalized linear model (development time) are shown in relation to relative contribution of EAA (classified using a Bayesian mixing model) from bacteria, fungi and plant (see Fig 7).

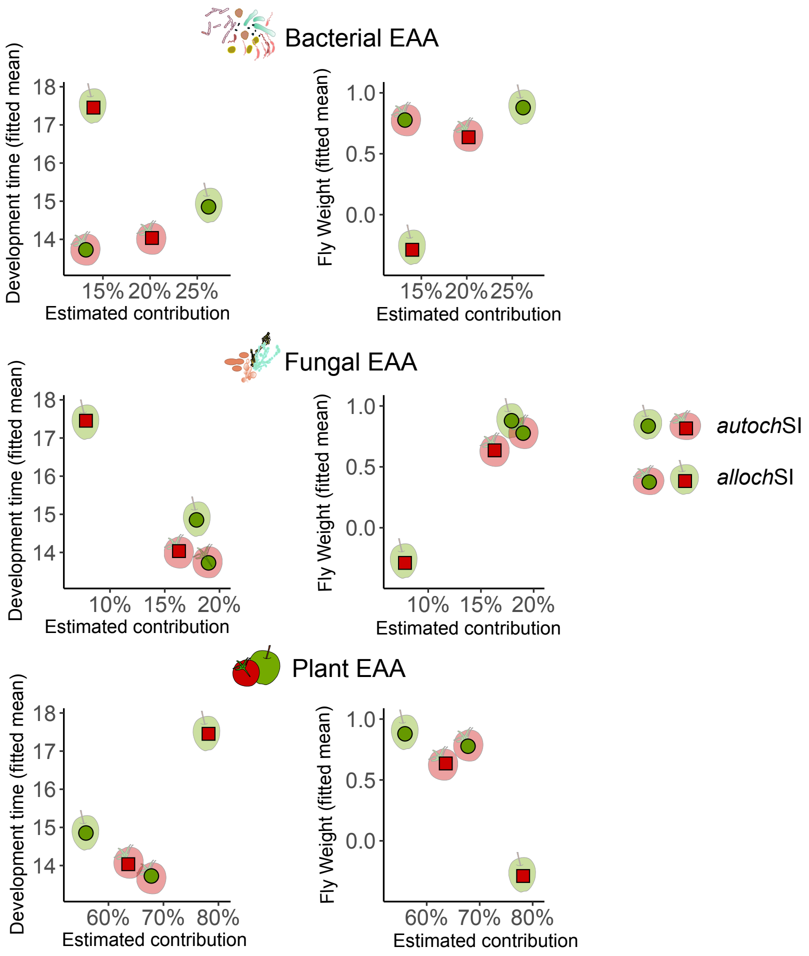

**Table S3. pH measurements of apple and tomato substrates**

Substrates were prepared from organic apples (Hollsteiner Cox) and organic vine tomatoes as described in the method section. They were homogenized and viscosity was adjusted reducing the water content of tomato substrate through sedimentation. Substrates were filled in 100 mL Schott bottles, two bottles for one substrate. One bottle of each substrate was heat sterilized. Bottles were left to cool to ambient temperature (ca. 20°C) before measurement on a Mettler Toledo Seven Direct SD20 pH-meter (d = 0.15 pH). Measurements were taken at two depths of the substrate: on the surface, covering only the probe in the substrate and in the substrate lowering the probe 0.5 cm into the substrate.

| Heat sterilized | Tomato substrate | Apple substrate |
|---|---|---|
| Surface measurement | 4.29 | 2.8 |
| Surface measurement | 4.31 | 2.64 |
| Surface measurement | 4.4 | 3.11 |
| Surface mean | 4.33 | 2.85 |
| 0.5 cm depth measurement | 3.91 | 3.16 |
| 0.5 cm depth measurement | 3.64 | 3.16 |
| 0.5 cm depth measurement | 3.98 | 3.18 |
| 0.5 cm depth mean | 3.84 | 3.17 |
| Fresh | Tomato substrate | Apple substrate |
| Surface measurement | 4.11 | 3.13 |
| Surface measurement | 4.14 | 3.11 |
| Surface measurement | 4.03 | 3.01 |
| Surface mean | 4.09 | 3.08 |
| 0.5 cm depth measurement | 4.16 | 3.19 |
| 0.5 cm depth measurement | 4.12 | 3.16 |
| 0.5 cm depth measurement | 4.1 | 3.18 |
| 0.5 cm depth mean | 4.13 | 3.18 |
